# Weekend light shifts evoke persistent *Drosophila* circadian neural network desynchrony

**DOI:** 10.1101/2019.12.31.891507

**Authors:** Ceazar Nave, Logan Roberts, Patrick Hwu, Jerson D. Estrella, Thanh C Vo, Thanh H Nguyen, Nicholas Pervolarakis, Paul J. Shaw, Tanya L. Leise, Todd C. Holmes

## Abstract

Billions of people subject themselves to phase-shifting light signals on a weekly basis by remaining active later at night and sleeping in later on weekends relative to weekday for up to a 3hr weekend light shift (WLS). Unnatural light signals disrupt circadian rhythms and physiology and behavior. Real-time light responses of mammalian suprachiasmatic nucleus are unmeasurable at single cell resolution. We compared *Drosophila* whole-circadian circuit responses between unshifted daytime/nighttime schedule and a 3hr WLS schedule at the single-cell resolution in cultured adult *Drosophila* brains using real-time bioluminescence imaging of the PERIOD protein for 11 days to determine how light shifts alter biological clock entrainment and stability. We find that circadian circuits show highly synchronous oscillations across all major circadian neuronal subgroups in unshifted light schedules. In contrast, circadian circuits exposed to a WLS schedule show significantly dampened oscillator synchrony and rhythmicity in most circadian neurons during, and after exposure. The WLS schedule first desynchronizes lateral ventral neuron (LNv) oscillations and the LNv are the last to resynchronize upon returning to a simulated weekday schedule. Surprisingly, one circadian subgroup, the dorsal neuron group-3 (DN3s), robustly increase their within-group synchrony in response to WLS exposure. Intact adult flies exposed to the WLS schedule show post-WLS transient defects in sleep stability, learning, and memory. Our findings suggest that WLS schedules disrupt circuit-wide circadian neuronal oscillator synchrony for much of the week, thus leading to observed behavioral defects in sleep, learning, and memory.

**Significance Statement:** The circadian clock controls numerous aspects of daily animal physiology, metabolism and behavior. Shift work in humans is harmful. Our understanding of circadian circuit-level oscillations stem from *ex vivo* imaging of mammalian suprachiasmatic nucleus (SCN) brain slices. However, our knowledge is limited to investigations without direct interrogation of phase-shifting light signals. We measured circuit-level circadian responses to a WLS protocol in light sensitive *ex vivo Drosophila* whole-brain preparation and find robust sub-circuit-specific oscillator desynchrony/resynchrony responses to light. These circuit-level behaviors correspond to our observed functional defects in learning and memory, and sleep pattern disruption *in vivo*. Our results reflect that WLS cause circadian-circuit desynchronization and correlate with disrupted cognitive and sleep performance.

## Introduction

Billions of individuals across the world subject themselves to phase-advancing light shifts on Monday morning after staying up later on weekends starting on Friday with phase-delaying light signals that persists throughout the duration of the weekend. Disruptions to circadian rhythmicity is linked to serious physiological illnesses such as heart diseases, diabetes, and cancers (Moore-Ede et al., 1983; Hastings et al., 2003; Scheer et al., 2009; Papagiannakopoulos et al., 2016). However, the effects of acute light shifts, like WLS, at the circadian-circuit level are not known.

PERIOD (PER) protein cycling imaging in suprachiasmatic nucleus (SCN) provides detailed functional data on the central neural circuits that govern circadian rhythms. Circuit PER cycling also forms the basis for interpreting the linkage between the timing of clock cycling and circadian physiological outputs in mammals (Welsh et al., 1995; Hamada et al., 2001; Yamaguchi et al., 2003; Evans et al., 2013; Azzi et al., 2017). Longitudinal optical or electrical recording of large numbers of neurons in SCN slices shows that free-running between-oscillator phases are complex and relatively desynchronized (Quintero et al., 2003; Schaap et al., 2003; Yamaguchi et al., 2003). Mammalian SCN ex vivo slices can be imaged at high spatial-temporal resolution, but their direct responses to environmental light signals cannot be studied due to the absence of retino-hypothalamic pathway input into the SCN in *ex vivo* preparations (Welsh et al., 1995; Moga and Moore, 1997; Hamada et al., 2001; Evans et al., 2013). This limitation prevents studies of how light shifts affect the neural circuitry that govern circadian rhythms.

The primary light input mechanism for the fly circadian neural circuit is via the blue-light sensitive photoreceptor CRYPTOCHROME (CRY) expressed in roughly half of the fly circadian neurons (Stanewsky et al., 1998; Yoshii et al., 2008; Fogle et al., 2011). A secondary and broad spectrally activated photoreceptor Rhodopsin 7 (Rh7) is also expressed in *Drosophila* circadian brain neurons and in photoreceptors (Kistenpfennig et al., 2017; Ni et al., 2017; Sakai et al., 2017). External opsin-based photoreceptors provide redundant photic input (Helfrich-Förster et al., 2001; Li et al., 2018). We took advantage of this feature of cell autonomous photoreceptors in many central brain circadian neurons to develop an imaging system that measures bioluminescence PER oscillation at single-cell resolution over many days in a long-term whole brain culture system (Roberts et al., 2015, 2016). Bioluminescence imaging of highly light-sensitive circadian neurons avoids all possibility of circuit perturbation by light contamination by fluorescence excitation (Hege et al., 1997; Roberts et al., 2015, 2016).

Our earlier work reveals circadian circuit-wide response of PER cycling to a single phase advancing light pulse consists of systematic desynchrony followed by resynchrony of PER cycling that varies between the different neuronal subclasses of the circuit (Roberts et al., 2015, 2016). The *Drosophila* whole brain culture responds to light-cued phase shifts throughout the circadian neural circuit in a very robust and reproducible fashion that appears to be identical to light cued phase shifts *in vivo*. This is shown by comparison of whole brain PER bioluminescence cycling with anti-PER immunocytochemistry prepared from whole flies exposed to light shifts *in vivo* (Zerr et al., 1990; Roberts et al., 2015, 2016). Here, we use *Drosophila* brains to study neural circuit response to WLS whose timing resembles the weekend/weekday light shifts experienced by many humans. Both mammalian and fly rhythms rely on circadian pacemaker circuit networks coupled by peptide and small molecule neurotransmitters (Renn et al., 1999; Maywood et al., 2006; Johard et al., 2009; Shafer and Yao, 2014; Jones et al., 2018). Motivated by the many functional similarities between mammalian and fly circadian circuitry, we investigated the effects of light shifts on circadian rhythmicity using *Drosophila*.

## Materials and Methods

### Behavioral Analysis of Day-Night Entrainment

Trikinetics *Drosophila* Activity Monitor (DAM) system was employed to record the locomotor activity of adult wild-type (W1118[5905]) and transgenic XLG-Luc flies (Roberts et al., 2015, 2016). Individual flies were placed in 5mm Pyrex glass tubes with fly food on one end, and a cotton plug on the other. Each experiment was run with either 32 or 64 adult male flies. The fly-containing tubes are mounted in a DAM5 *Drosophila* Activity Monitor (TriKinetics) which records the number of infrared beam crossings over time, as a measure of activity. Flies are first entrained under standard 12hrs light: 12hrs dark (12:12LD) conditions for ≥3 days. Following entrainment, flies are then exposed to either the *LD Strobe* protocol that for each hour of “light” 15minLight:45minDark is repeated ever hour for 12 hours superimposed over a 12:12LD; Skeleton Photoperiod (SPP); or standard 12:12LD light protocols with consistent phases and a consistent (white light intensity: of 1.1 mW/cm^2^) for 8 days. The *LD Strobe* protocol is performed by dividing the 12hrs of daytime entrainment into 12 one-hour cycles of short, intermittent light-dark exposures followed by 12 hours of nighttime darkness. The 15-minute skeleton photoperiod protocol is performed by applying a short light pulse at the transition times of expected lights on (simulated dawn) and lights off (dusk) based on the previous standard entrainment. Initial optimization tests for *LD Strobe* and SPP protocols were determined with light pulse durations of either 5, 15, or 30 m using highest behavioral circadian rhythmicity under subsequent constant darkness (DD) conditions as comparison criteria. Following the 8 days of entrainment by either 12:12LD, LD Strobe, or SPP, we examined the free-running circadian activity of the flies for ≥3 days under DD.

### Quantification of Locomotor Activity

FaasX (M. Boudinot and F. Rouyer, Centre National de la Recherche Scientifique) was used for analysis of locomotor activity recorded by the automated Trikinetics *Drosophila* Activity Monitor system. Cycle-P was utilized to quantify period length, amplitude and rhythmicity using 15-minute bins of individual fly locomotor activity. Individual fly rhythmicity is defined rhythmic based on chi-square periodogram analysis with the following criteria (high frequency filter on): power ≥40, width ≥4 hours and period length of 24 ±8 hours. Double-plotted actogram graphs were generated by the software ClockLab (Actimetrics) showing normalized activity over 1-minute intervals.

### Bioluminescence Imaging

Custom bioluminescence set up is designed and built by Logan Roberts with David Callard, and Jeff Stepkowski (Stanford Photonics) and Todd Holmes. Bioluminescence set up includes custom light filters, LED light set up by Prizmatix, a retooled and light-tight black box, and custom temperature control maze. Bioluminescence imaging is performed using adult, male XLG-Per-Luc transgenic fly brains (line provided by Ralf Stanewsky, University of Münster, Germany, (as descibed in Veleri et al., 2003). XLG-Per-Luc flies are first entrained to ≥3 days of 12:12LD entrainment before dissection. Six whole fly brain explants are dissected and cultured on a single insert per experiment using a modified version of a previously described protocol (Roberts et al., 2015). The cultured brains are mounted on a stage (Applied Scientific Instrumentation) with automated XYZ movement controlled by the software Piper. The stage is connected to an upright Axio Observer.Z1 Microscope (Zeiss) set in a custom light-tight incubator (designed by Alec Davidson, Morehouse School of Medicine, GA) with temperature maintained at 25°C ±0.5°C. Bioluminescence from the cultured whole brains is collected by a Zeiss 5x (NA=0.25) objective and transmitted directly to a MEGA-10Z cooled intensified CCD camera (Stanford Photonics) mounted on the bottom port of the microscope. The XY position of the samples is manually set using bright-field illumination. The optimal z-plane of focus for bioluminescence imaging is obtained by performing 10 Z-steps at 40-50 µm intervals with 5-10-minute exposures. Experimental bioluminescence imaging of the samples is obtained with 15-minute exposures at 30 fps for ≥11 days of recording at single-cell resolution during the hourly dark phase of the LD strobe protocol. Light exposure and entrainment are performed using an *LD strobe* protocol with the 12 hours of daytime entrainment divided into 12 consecutive cycles of a 15-minute light pulse and 45 minutes of darkness, followed by 12 hours of constant darkness (hereby referred to as “15L45D/*LD Strobe*”). Images are collected by Piper (Stanford Photonics) and averaged into 45-minute bins by ImageJ before using MetaMorph (Molecular Devices, Sunnyvale, CA), Microsoft Excel and custom MATLAB scripts to measure circadian parameters of bioluminescence cycling with single cell resolution. Only experiments with all six brain explants still healthy, contamination-free, adhering to the insert substrate, and exhibiting bioluminescence for ≥11 days are used for analysis.

### Simulating day-night entrainment and weekend light shifts *ex vivo*

To establish baseline measurements of day-night entrainment, one group (referred to as the control group) consists of whole brain explants exposed to the 15L45D *LD Strobe* schedule that simulates 12:12LD entrainment for 8 days with no phase shifts followed by ±3 days of constant darkness (DD). Stable white light exposure (30µW/cm^2^, as performed in our previous published work (Roberts et al., 2015, 2016) using a mic-LED (Prizmatix)) is set to provide a stable light intensity with automated timing set via TTL input from Piper (Stanford Photonics). During intervals of light exposures, the CCD camera is protected by a mechanical shutter controlled via TTL input from Piper to allow for semi-continuous imaging. For samples exposed to a WLS protocol (referred to as WLS), the first three recorded “weekdays” (all with the same phase for a simulated “Wednesday” to “Friday”) have parallel phases with the control group. This is followed by a 3-hour phase delay on the evening of the third recorded day (simulated “Friday night”) followed by two “weekend” days but with no phase shift (simulated “Saturday” through “Sunday”). Whole brain explants are exposed to a phase advance of three hours on the morning of the sixth day of recording (simulated “Monday” morning) with no phase shifts for the following simulated weekdays (“Monday” through “Wednesday”). Finally, explants are placed in constant darkness (DD) for ≥3 days. Three-hour phase shifts were used because they correspond with social behaviors commonly observed in the general populace regarding weekend light shifts and have been linked to negative health effects. LED light exposure and brain imaging are automated via TTL input through the Piper software provided by Stanford Photonics (control group=208 players, WLS group=214 players).

### Processing of Bioluminescence Images

Cosmic rays are removed in real-time using the Piper cosmic ray filter set to discriminate the sum of all pixel values above 800 and reject frames that are >3 standard deviations over the running average (run over 30 frames). ImageJ is used to generate images with bioluminescence images averaged over 45-minute intervals. These images were then further processed using MetaMorph as described in previously work (Roberts et al., 2015, 2016). Briefly, noise from dark current and cosmic rays were removed by using a running minimum algorithm to generate new images constructed from pairs of sequential images using the minimum values of each pixel from the two images. MetaMorph was used to generate a stack of images for each experiment with average luminescence intensity over time measured for regions of interest (ROIs) that were manually defined based on a previous protocol (Roberts et al., 2015). ROIs were classified into canonical circadian neuron groups (colored-coded: red = s-LNv, yellow = l-LNv, orange = LNd, blue = DN1, green = DN3) based on consistent and classically recognized anatomical locations. Raw bioluminescence data were then processed by Microsoft Excel was used to adjust for background noise and convert raw luminescence over time to photons-per-minute as previously described (Roberts et al., 2015, 2016). Circadian parameters were analyzed for 11-day recordings using modified versions of previously described MATLAB scripts with the first 12hrs excluded due to initially high amplitude and highly variable bioluminescence following dissection and addition of luciferin (Roberts et al., 2015). Between circadian neuronal cell group variable bioluminescence persists for several days after dissection. These records are retained to show re-emergence of highly synchronized between circadian cell group rhythms after several days in culture. This is in strong agreement with anti-PER immunocytochemical “snapshots” of highly synchronous *in vivo* fly brain PER cycling measured in flies maintained in LD over 24 hours (Zerr et al., 1990).

### Quantification of Circadian Oscillator Dynamics

Custom MATLAB scripts (version 8.2) were employed to analyze real-time bioluminescence recordings for quantification of order parameter, goodness-of-sine-fit, amplitude, period and phase. The order parameter ‘R’ was used to quantify the synchrony of phase, period and waveform for each circadian neuron subgroup and for ‘all cells’ (summed from all subgroups). Statistical significance was determined by using a bootstrap procedure with the size of each bootstrap sample equivalent to the original number of cells in each data set. Each bootstrap procedure was repeated 4,000 times (except 10,000 for the ‘all cells’ group) to provide 95% and 99% confidence intervals for the difference in R (RWLS-RC) between cells exposed to WLS and cells in control conditions with no phase shifts with the null hypothesis that there is no difference between conditions. Discrete wavelet transform (DWT) was used in combination with sine-fit estimates of 2-day sliding windows to provide circadian measures of rhythmicity, period, amplitude and phase. Oscillator rhythmicity was determined as the percentage of variance accounted for by fitting a sine wave to the time series (goodness-of-sine-fit). Oscillators were deemed “reliably rhythmic” if their period was 24 ±8 hours, amplitude was the noise amplitude (mean amplitude of the DWT component associated with periods shorter than 4hrs), and their goodness-of-sine-fit measure was ≥0.82 as described in (Roberts et al., 2015). Circular phase plots were generated using the midpoint of 2-day sliding windows with an inner circle showing the alpha=0.05 threshold for the resultant vector for the Rayleigh test. Circular plots were standardized with ZT0 set equal to the overall mean phase of “all cells” in the control condition on day 3 when entrainment is most stable. The absence of an oscillator’s plots at certain time points indicates that the oscillator’s rhythmicity did not meet the criteria for reliably rhythmic cells and was thus too dampened to reliably measure.

### Non-linear Embedded Phase Estimates Used to Generate Phase Ensemble Animations and Validate Sine-Fit Estimates

A time delay embedding protocol was used to confirm that circadian parameter s was reliably measured using sine-fits of wavelet detrended time series. Phase estimates were determined by the polar angle of time series that were embedded in a higher dimension via a 6-hour lag resulting in oscillations circling the origin. Phase plots generated using this nonlinear embedded phase analysis confirmed the same patterns of oscillator dynamics observed in plots generated using sine-fit calculations. Non-linear embedded phase estimates were also used to generate phase ensemble animations as previously described (Roberts et al., 2016). Dynamic changes in phase and amplitude were displayed at 3 levels of resolution: whole circadian neuron network (Movie S2), individual neuronal subgroups (Movie S3), and single neuron oscillators (Movie S4). In Supplemental Movies 2 and 3, the polar angle of each disk represents the relative phase shift in control and weekend light shift conditions. The size and proximal distance of the disks from the center of the circle represents amplitude (i.e. damping is indicated by a shrinking disk drifting towards the center). In Supplemental Movie 4, the polar angle of each disk indicates the phases of individual neuron oscillators while amplitude is again reflected by the size of the disks and proximal distance from the center.

### Fly Sleep Quantification

Fly activity data was binned into 60-minute time sections. Following binning, activity data was translated to run length encoding and any run of zero activity for five or greater minutes was scored as sleep. Each fly’s total amount of sleep per bin was totaled, and the resultant matrix contained the total amount of sleep by fly per 60-minute increments for the length of the experiment. Flies that died during the course of the experiment registered very long strings of zero activity and were manually removed to prevent over counting sleep amounts. Students two-sided t-test was used to compare differences between bins of experimental groups, with a p-value <0.05 considered to be significant.

### Learning and Memory Assay

Flies were evaluated for the effects of the weekend light shift on both sleep and short-term memory (Seugnet et al., 2009). ∼6-day old male Canton-S (Cs) flies were used to assess short-term memory (STM) using the Aversive Phototaxic Suppression (APS) assay. Prior to being tested for STM, flies are examined to determine if they exhibit normal photosensitivity and quinine photosensitivity. This step is used to ensure that the changes to sensory thresholds are true changes in associative learning. Photosensitivity is evaluated using a T-maze with one lightened and darkened chambers that appear equal on either side. Flies must make photopositive choices to be considered for post-WLS evaluation of STM. Quinine sensitivity index (QSI) is achieved by determining the duration a fly stay on a side of a T-maze without quinine, as opposed to one side with the aversive stimuli, during a 5-minute period (Seugnet et al., 2009). The learning test to evaluate STM used the APS assay. Flies were subjected to WLS schedule used for bioluminescence and behavior experiments. Flies were tested on the subjective “Tuesday” of the WLS schedule, two days after 3hr phase shifts experienced during the weekend. Flies are individually placed in a T-maze and allowed to choose between a lighted or darkened chamber for over 16 trials. Flies that do not display phototaxis during the first block of four trials are excluded from further trials. During the 16 trials, flies learn to avoid the lighted chamber paired with aversive stimulus (Seugnet et al., 2009). The performance index is calculated as the percentage of the times the fly choses the dark vial during the last four trials of the 16-trial test. STM is defined as selecting the dark vial on two or more occasions during the final four tests.

## Results

### Development of LD Strobe to Simulate Day-Night Entrainment

Light is the primary environmental cue for circadian entrainment (Hall, 2005). Previous studies show that *Drosophila* brains are directly light sensitive due to the expression of cell-autonomous, short-wavelength, light-sensitive photoreceptors. Light-sensitive components CRYPTOCHROME (CRY) and Rhodopsin-7 (Rh7) are expressed in select circadian neuronal subsets and are distributed throughout the fly brain (Stanewsky et al., 1998; Veleri et al., 2003; Benito et al., 2008; Yoshii et al., 2008; Fogle et al., 2011; Ni et al., 2017). The sensitivity of the fly brain to light input enables the study of physiological photic entrainment using real-time bioluminescence recordings of entire cultured brains (Roberts et al., 2015, 2016). In addition to long durations of light, circadian cycles can be entrained using short pulses of light, referred to as “skeleton photoperiods (SPP)” (Pittendrigh and Minis, 1964; Pittendrigh and Daan, 1976). In SPP, light pulses flank the beginning and end of the simulated daytime, which is then followed by long periods of complete darkness (DD) that simulate nighttime. The ability for entrainment using short periods of light at the correct time points makes suggests that obtaining bioluminescence images during simulated daytime is possible. However, circadian locomotor behaviors in DD that follow skeleton photoperiods are weak and differ significantly from the robust circadian behaviors seen in DD following standard 12h:12h light:dark (LD) cycles (Fig. S1).

As circadian behavior in DD reflects the activity of the free-running clock, we established the criteria that DD behavior following imaging conditions must show no statistically significant differences compared to normal LD cycles, as DD behavior reports the state of the circadian clock circuit. For the purposes of entire CNS circadian circuit imaging, we developed a novel entrainment protocol we refer to as *LD Strobe* (Fig. S1, also see methods). *LD Strobe* consists of 15-minute periods of light followed by 45-minute bouts of darkness each hour during the 12-hour “day”, then 12 hours of darkness during the 12-hour “night.” DD following *LD Strobe* is indistinguishable from that following standard LD (Fig. S1). Thus, *LD Strobe* effectively simulates daytime during 12 hours of alternating periods of light and dark, where dark periods provide the opportunity to capture circadian circuit bioluminescence.

### Exposure to weekend light shifts dampen circadian circuit-wide rhythmicity and synchrony

With the ability to study bioluminescence during simulated daytimes, we examined the circadian circuit-wide dynamic response at the single-neuron resolution to compare unshifted to shifted light schedules. For 11 days, we obtained bioluminescence imaging of cultured adult *Drosophila* brains exposed to the *LD Strobe* entrainment schedule. Under control (CTRL), unshifted LD conditions, we find a high level of multi-day synchrony (8 days) between all major circadian neuron subgroups (Fig. 1A) along with robust rhythmicity, high amplitude oscillations, and ∼24hr. After 8 days of unshifted LD, we challenged the free-running clock by placing brains in constant darkness (DD). Despite an immediate loss in clock cycling amplitude between canonical circadian subgroups in DD, they remain synchronized as cycling amplitude dampens (Fig. 1A, gray shade).

**Figure 1.**
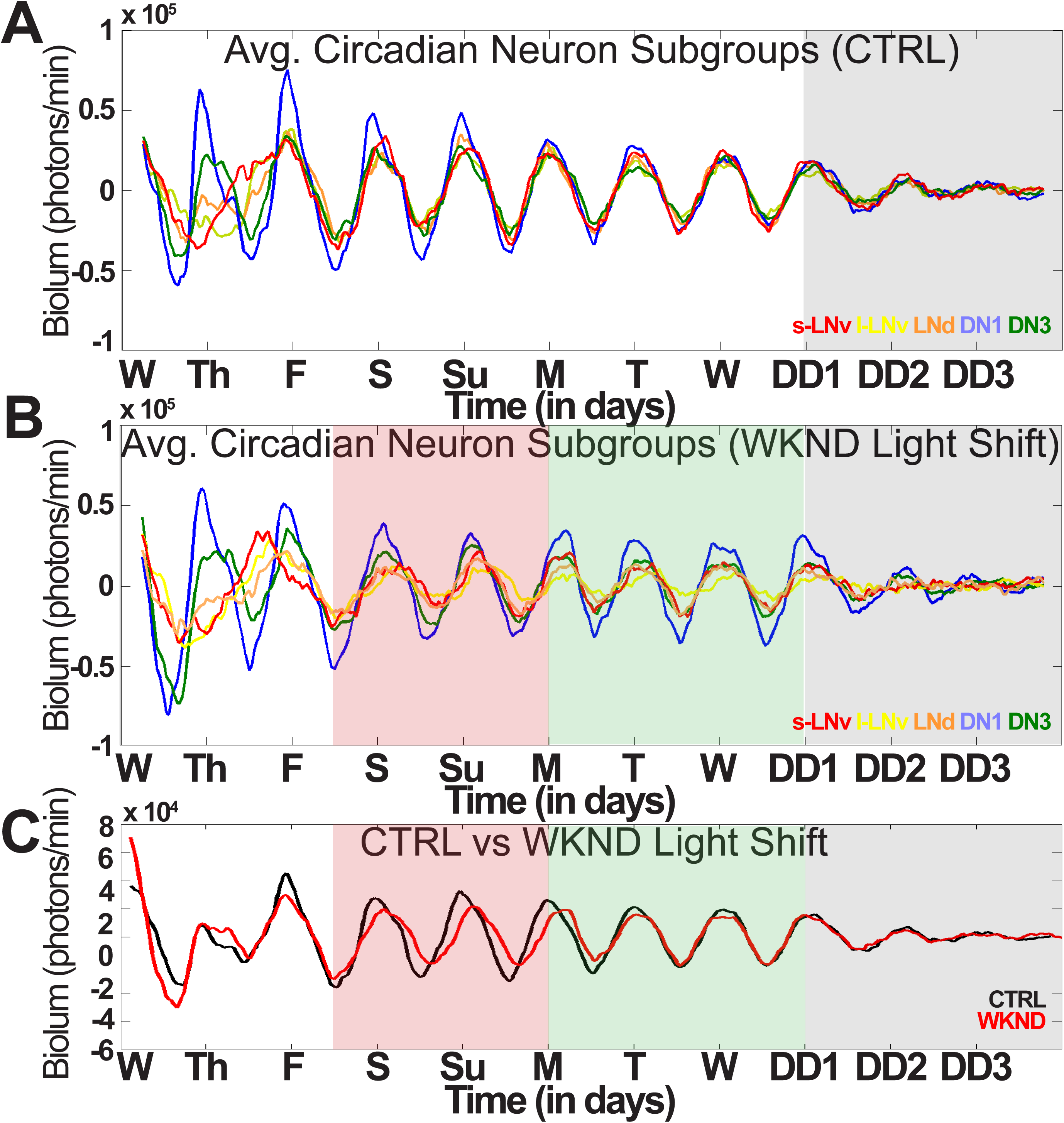
Weekend light shifts dampen circuit-wide rhythmicity and synchrony. 11-day bioluminescence recordings of cultured *Drosophila* brains reported by PER (n=6). (A) Control LD conditions simulate standard 12L12D entrainment spanning one week without phase shifts (sLNV n=18, l-LNv n=19, LNd n=18, DN1 n=27, DN3 n=18) followed by DD (gray shade). (B) WLS conditions subject brain cultures to one simulated weekend with 3hr phase delay on Friday (red shade), 3hr phase advance on Monday (green shade), followed by three days of DD (gray shade) (sLNV n=17, l-LNv n=17, LNd n=15, DN1 n=28, DN3 n=30). (C) Averaged bioluminescence tracings in control LD (black) or WLS conditions (red).

To simulate WLS, we performed light phase shifts by initiating a 3hr phase delay, simulating “staying up late on Friday.” We retained this 3hr phase delayed schedule for two days simulating “sleep in late, stay up late” followed by a 3hr phase advance to simulate “Monday morning.” Similar weekday/weekend schedules, as previously described, are chronically experienced by much of the human populace worldwide. We find that *Drosophila* whole-brain explants exposed to WLS schedules show reduced synchrony between and within canonical circadian neuron subgroups during (Fig. 1B, red shade) and after simulated weekend phase shifts (Fig. 1B, green shade). Furthermore, circadian neuron subgroups following WLS show an immediate loss in rhythmicity and synchrony during the transition into DD (Fig. 1B, green shade), revealing long-lasting circadian neural circuit perturbation three days after the last phase shift that simulates one weekend. This contrasts with the high level of synchrony seen in DD for unshifted CTRL (Fig. 1A, gray shade). Averaged circuit-wide cycling (Fig. 1C) is compared between unshifted CTRL (black) and 3hr WLS (red), indicating average oscillator phase recovery does not occur until several days post-shift.

The striking differences in oscillator rhythmicity and phase coherence found in CTRL LD (Movie S1, left) are observed when compared to brains exposed to WLS (Movie S1, right). Phase ensemble animations aid to visualize oscillator dynamics comparing brains in CTRL (left) and WLS (right) schedules averaged as a whole throughout the duration of the experiment (Movie S2), separated into canonical circadian neuron subgroups (Movie S3), and at single-cell level (Movie S4). Differences in inter-subgroup dynamics in CTRL and WLS conditions led us to investigate circadian cycling of individual circadian neuron subgroups at single-cell resolution.

### Individual circadian neuron subgroups exhibit distinct dynamics of activity under WLS conditions

Under unshifted CTRL LD, circadian neuron subgroups exhibit distinct signatures in rhythmicity, phase coherence and amplitude throughout the duration of entrainment (Fig. 2A). Small-lateral ventral neurons (sLNvs) show the highest degree of synchrony over the course of CTRL LD (Fig. 2A, first column). Dorsal neurons-1 (DN1) exhibit the highest amplitude oscillations (Fig. 2A, fourth column), and the DN3s exhibit the greatest variability in oscillator amplitude and synchrony in CTRL LD (Fig. 2A, fifth column). Quantitatively, sLNvs and lateral-dorsal neurons (LNds) maintain the highest level of synchrony in CTRL conditions (Fig. 2C, red and orange), whereas DN3s exhibit the lowest (Fig. 2C, green). In DD, following CTRL LD, all neuronal subgroups exhibit a clear and immediate decrease in amplitude (Fig. 2A, blue trace, gray shade).

**Figure 2.**
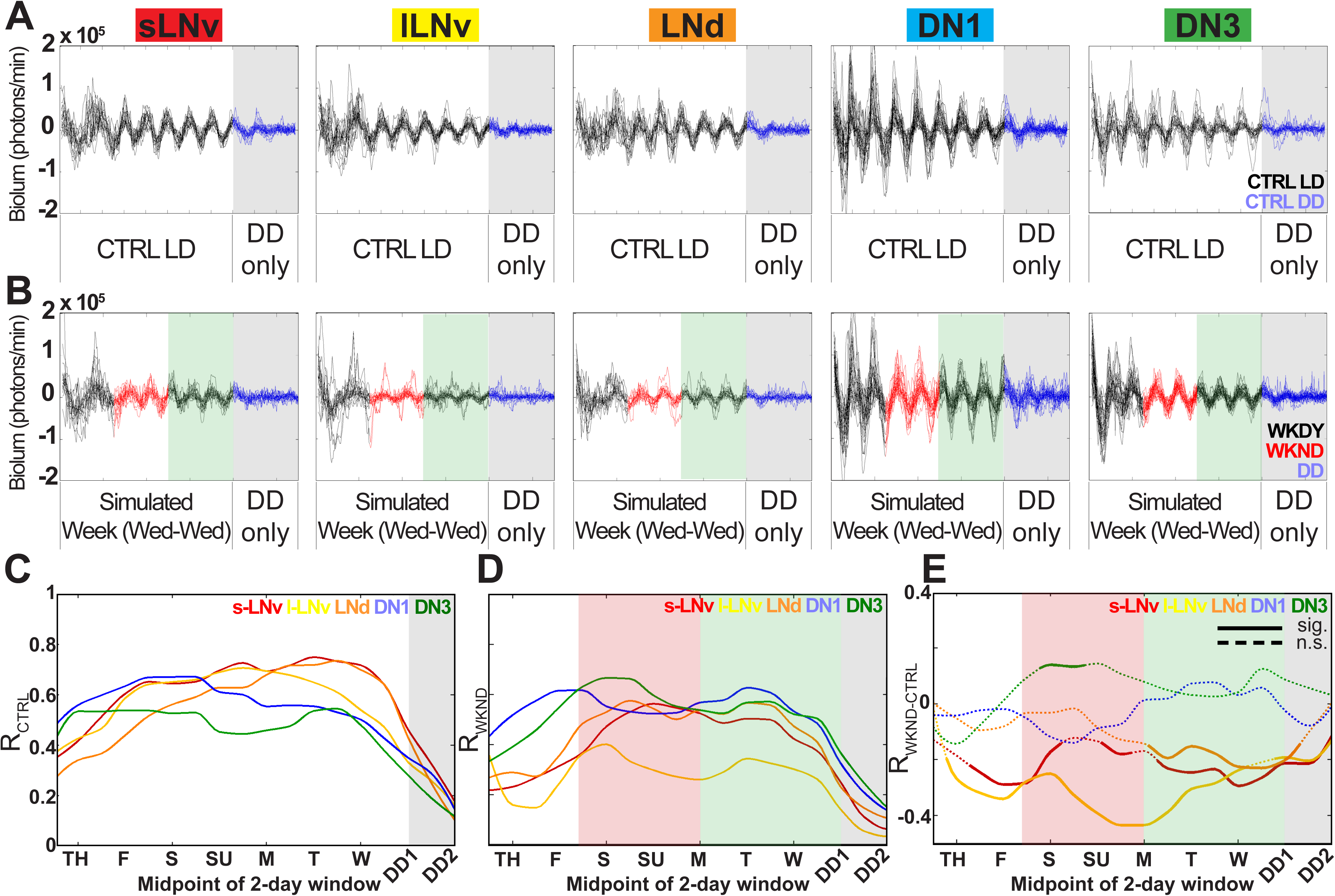
Circadian neuron subgroups exhibit significantly distinct dynamics of activity in response to WLS conditions. (A) Control LD conditions for each circadian neuron subgroup, each line represents an individual cell. Brains were subjected to DD (blue trace) after a simulated week. (B) WLS conditions for each circadian neuron subgroup, each line represents an individual cell in pre-WLS (black trace), WLS (red trace), post-WLS (green shade), then three days of DD (blue trace, gray shade). Calculated dynamic changes in order parameter R as a measure of synchrony for individual neuron subgroups in control LD (C) or WLS (D) conditions. (E) Statistical comparisons of inter-subgroup synchrony using 2-day sliding windows. Dotted lines indicate no significant changes between control LD and WLS conditions, solid lines indicate significance outside the 95% confidence interval calculated using bootstrap analysis.

WLS disrupt rhythmicity and synchronization during and after shifts affecting all circadian neuron subgroups except for DN3s (Fig. 2B) which significantly tighten their amplitude and phase coherence (Fig. 2D, E). Interestingly, DN3s are the only cells that become significantly more synchronized in response to WLS returning to a “weekday” schedule (Fig. 2B, fifth column, green shade, and Fig. 2E, green trace). Small- and large-LNvs show significantly lower synchrony in response to WLS relative to unshifted CTRL LD (Fig. 2E, red and yellow traces). Conversely, LNds, DN1s, and DN3s maintain robust synchrony and amplitude in response to WLS (Fig. 2E, orange, blue and green traces). Despite being cell autonomously light-blind by not expressing CRY or Rh7 like other subgroups (Benito et al., 2008; Yoshii et al., 2008; Ni et al., 2017), DN3s significantly increase inter-group oscillator synchrony as a response to WLS relative to unshifted CTRL LD (Fig. 2B, fifth column, Fig. 2E, green trace).

Oscillator phase remarkably stabilizes within and between all circadian oscillator subgroups in CTRL LD, particularly for sLNVs which maintain day-to-day average phase close to CT0 (Fig. 3A and C, see 3D for detail of circular plot). In DD, only sLNVs show similarly stable day-to-day average phases close to CT0 (Roberts et al., 2016), consistent with suggested roles as the dominant circadian subgroup in complete darkness. At intervals following a “Friday” WLS phase delay, all oscillator groups either immediately (sLNVs, lLNvs) or gradually (DN1s) shift to a more negative average phase values relative to CT0 seen in CTRL LD (Fig. 3B). Moreover, all circadian neurons, except for sLNVs, gradually return to phase values indistinguishable from CTRL LD two to three days following the “Monday” 3hr phase advance (Fig. 2E, Fig. 3B, and Fig. S3A, B). This is a markedly different response observed following a single15-minute light pulse that evokes a 2hr phase advance in the midst of DD conditions, leading to large, positive phase shifts for most circadian subgroups (Roberts et al., 2015, 2016).

**Figure 3.**
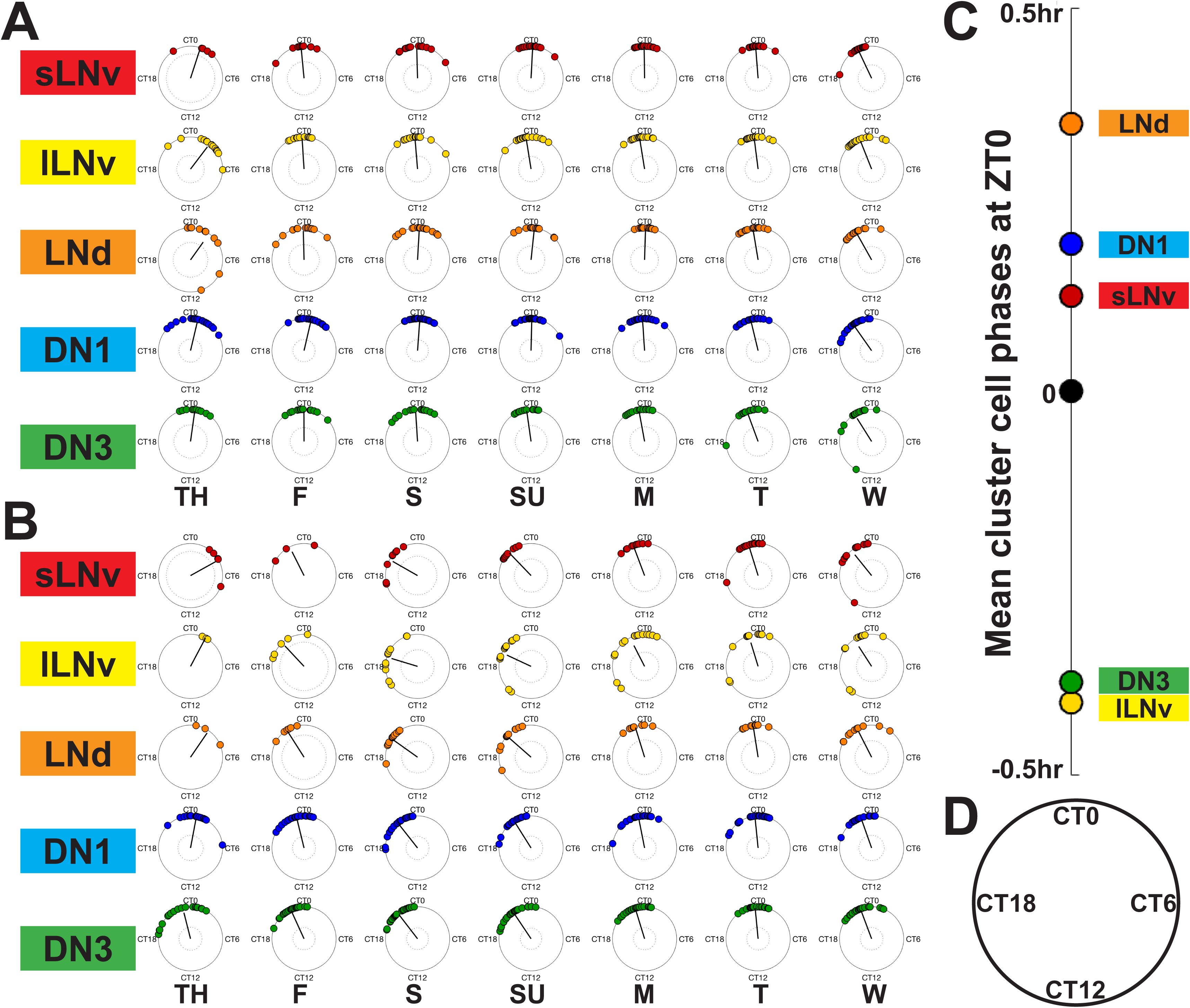
Circadian neuron subgroups exhibit stable dynamics in control LD but destabilize during WLS and requires days to recover post-shift. (A) Circular phase plots show all neuronal subgroups have high phase coherence within and between subgroups in control LD for one simulated week. (B) Neuronal subgroups dynamically shift phases during and after weekend phase delays, and phase advancing light shifts with greater phase dispersal. Dotted circles in (A) and (B) indicate the α=0.005 threshold for the Rayleigh test on the resultant (straight line); tests are no run if sample size is less than four. (C) Average phase values for individual subgroups in control LD. (D) General schematic of circular phase plot to show 24hr cycle.

### Exposure to WLS leads to sleep disruption, and defects in learning and memory

Due to long-lasting post-shift changes in oscillator ensemble activity in circadian neurons following WLS, we measured correlative behavioral outputs under the similar light shift protocols *in vivo*. We exposed whole, intact flies to CTRL LD and WLS schedules used in while-brain imaging while measuring sleep (Hendricks et al., 2000; Tononi, 2000). Sleep is stable for unshifted CTRL LD (Fig. 4A) as shown by consistent amplitudes (x-axis) and robust waveform (y-axis) across multiple days (z-axis). Flies exposed to WLS have significantly disrupted sleep patterns only during and after phase shifts (Fig. 4B, red dots indicate significant hourly difference in sleep compared to CTRL LD). Hourly sleep differences between CTRL LD and WLS groups persist up to six days following the “Monday” phase advance into DD (Fig. 4B, gray shade). Though hour-by-hour sleep amounts differ between CTRL LD and WLS groups during days after the phase advancing light shift, the total sleep amount over time does not differ—indicating the eventual effect of sleep homeostasis. Decreased cognitive performance, including learning and memory, is linked to circadian dysregulation and sleep low (Stickgold et al., 2001; Donlea et al., 2011). We tested how WLS affects learning and memory using the Aversive Phototaxic Suppression (APS) assay (Seugnet et al., 2009) two days after the phase-advancing light shift. WLS flies show significant impairments in remembering where to avoid aversive stimuli (quinine) in the maze compared to flies in CTRL LD (Fig. 4D). This suggests that WLS exposure impairs learning and maintenance of short-term memory days following weekend phase shifts as shown by the inability for conditioning to avoid aversive stimuli, coinciding with persistent post-shift circadian circuit and sleep pattern impairment.

**Figure 4.**
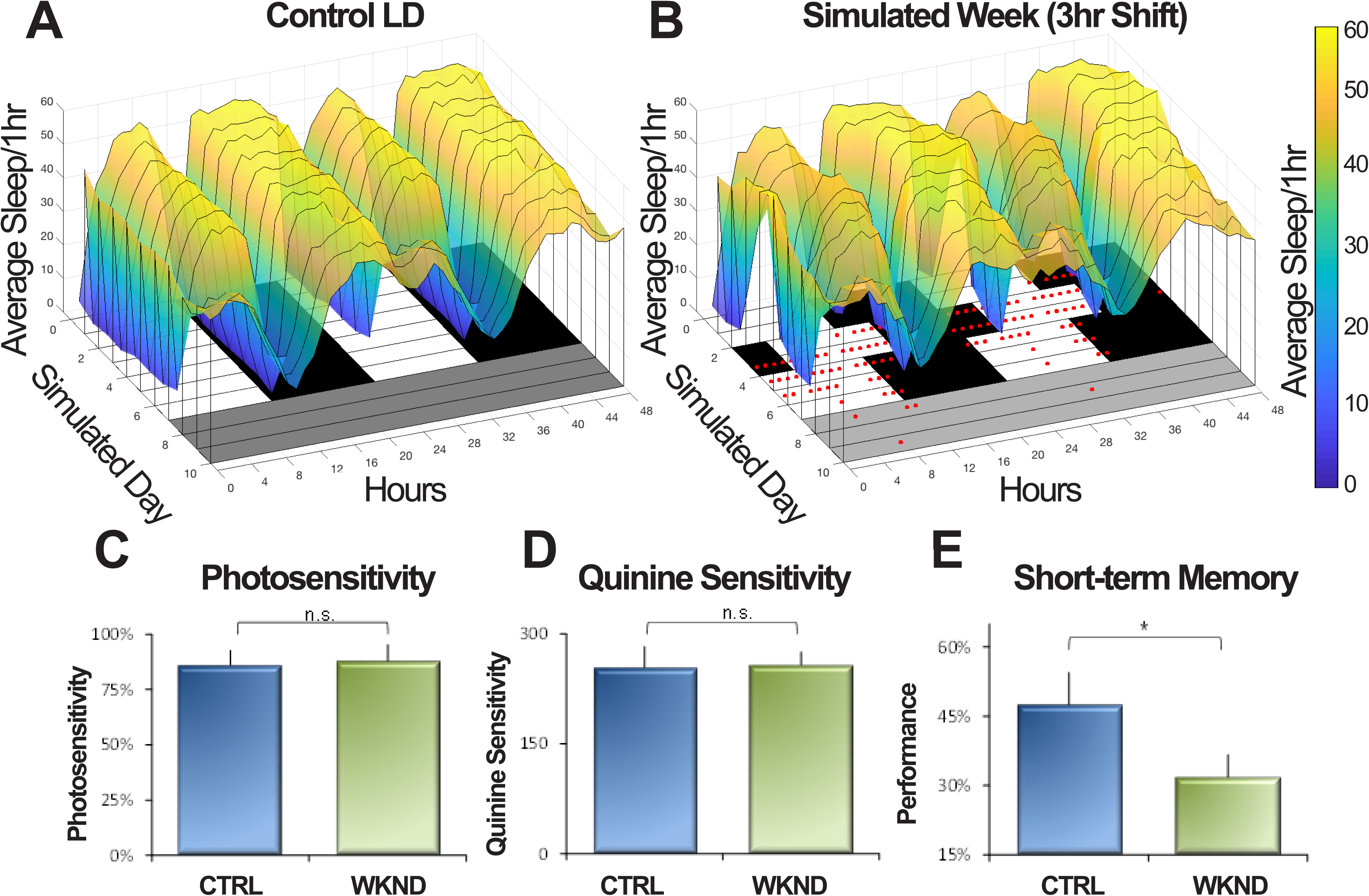
WLS disorder sleep and hinders learning and memory. Three-dimensional heat maps compare sleep profiles in control LD (A) and WLS conditions (B) as 1hr bins. Increased sleep amount is seen in yellow and lowest amount/absence of sleep in blue. Each row represents day (white bars) and night (black bars) spanning two simulated days (double plots); experiments last one simulated week followed by DD (gray shade). Red dots indicate significant differences in amount of average sleep/1hr bin between WLS and its respective control LD condition. (C) Photosensitivity assay comparison for flies in control LD or WLS conditions two days after phase advance (n=5/condition). (D) Control quinine sensitivity measurements for flies placed in control LD and WLS conditions two days after phase advance (n=5/condition). (E) Aversive Phototaxic Suppression (APS) assay two days after phase advance between control LD and WLS conditions (n=10-11 flies/condition). Significance for sleep differences was determined using a t-test at p<0.05.

## Discussion

Our whole-circuit imaging using *LD Strobe* provides longitudinal multi-day bioluminescence recordings of the entire circadian neural network at single-cell resolution. Moreover, under simulated LD cycles, we obtained detailed circuit responses to regular light: dark cycles and light shifts approximating weekday and weekend light shift conditions. We find coherent mean circadian network phase in response to alternating light pulses of *LD Strobe*. This indicates that properly timed Zeitgebers at the start and end of daytime hours, with standard night-time darkness are the most critical light input features for proper entrainment, consistent with earlier studies employing skeleton photoperiod protocols (Pittendrigh and Daan, 1976). In contrast, oscillators exhibit immediate damping in rhythmicity and synchrony when transitioning from LD to DD indicating that steady photoentrainment is critical for maintenance of robust oscillator synchrony and physiological rhythmicity (Roberts et al., 2015, 2016).

Different anatomically defined subgroups of circadian neuronal oscillators exhibit varying degrees of change in inter- and intra-subgroup synchrony and amplitude in response the WLS phase delay followed by a light phase advance days later. Single-cell analysis reveals that rhythmicity and synchrony in sLNvs, lLNvs, and LNds immediately dampens by initial “Friday” phase delay to mark the start of WLS. Taking the LD and DD data together, the sLNVs are the most stable of all circadian subgroups in the absence of light shifts but are most labile in response to changes in light entrainment evoking the “first out, last back in” phase destabilization (Roberts et al., 2016). In contrast, DN1s while showing light-induced phase shifts, maintain greater rhythmicity and synchrony in response to the phase delays. Surprisingly, the DN3s significantly increase their synchrony in response to phase delays in addition to the striking reduction in oscillator amplitude variance. LNds and the light blind DN3s immediately restore rhythmicity faster than other subgroups after exposure to a simulated weekend. Remarkably, the DN3s increase in phase synchrony and amplitude coherence during and after WLS entrainment. The DN3s could code for the initial time phase before light induced shift to a new phase, thus acting as a temporal placeholder. The LNds and DN3s may play a critical role in prompting the remaining circadian neural network into a new state of adaptation of the phase-shifted synchrony, consistent with earlier evidence indicating LNDs track phase-advance shifts more rapidly than other subgroups (Roberts et al., 2015, 2016).

Furthermore, light input from external photoreceptors, such as the compound eye, also modulate clock entrainment (Helfrich-Förster et al., 2001). Light from either input channel in the absence of the other (CRY/Rh7 versus external opsin expressing photoreceptors) yields identical behavioral light circadian phase shift responses (Helfrich-Förster, 2001; Ni et al., 2017), indicating these input channels are functionally redundant and the circadian circuit likely encodes their input in a similar if not identical fashion. Further, recent work shows electrophysiological light responses in circadian neurons using recording light stimulus parameters that are optimized for opsin activation in eyes but are insufficient in duration (and perhaps amplitude) for CRY activation (Li et al., 2018; see Baik et al., 2019 for detailed light parameters for CRY activation). As expected, we find that overall circuit entrainment by direct light is comparable whether the compound eye is present or is completely removed (Fig. S4). Additionally, cultured brains respond to the phase shifts of WLS schedules in a similar manner regardless of the presence or absence of compound eye in culture (Fig. S5). This confirms the functional redundancy of internal cell autonomous photoreceptors and external opsin-based photoreceptors.

The significant loss in amplitude and synchrony between oscillators following WLS relative to CTRL transitioning to DD suggests long-lasting residual circadian circuit instability is masked by light inputs into the neural circuit. Together, the data indicate long-lasting desynchrony of the circadian circuit during and after phase delays and advances of the WLS “weekend.” The circadian circuit is likely destabilized for the greater part of the week for individuals that shift every weekend as a matter of lifestyle. Considering the correlative defects in post-WLS sleep stability, learning, and memory, this poses the critical question of whether these defects are cumulative and can be detrimental over time. Phase shifts due to “jetlag” disrupt the timing of both arousal/wake and sleep. Circadian neurons functionally segregate to control arousal (sLNvs, lLNvs) versus sleep (DN1s) (Parisky et al., 2008; Shang et al., 2008; Sheeba et al., 2008; Guo et al., 2016). *In vivo* luciferase calcium monitoring at circadian neuronal subgroup spatial and temporal resolution confirms that the sLNv and lLNv intracellular calcium signaling exhibit biphasic morning and late day peaks corresponding to arousal while a subset DN1s coincide highest intracellular calcium levels with sleep (Guo et al., 2016, 2017). This innovative imaging approach yields robust records of circadian signal transduction occurring in these neurons in the absence of potential contaminating light excitation necessary for fluorescence imaging.

Circadian clocks regulate numerous aspects daily animal physiology and behavior. In humans, disruption of the circadian clock, and sleep loss, contribute to numerous diseases (Moore-Ede et al., 1983; Filipski et al., 2004; Scheer et al., 2009). Light is the primary environmental zeitgeber for circadian entrainment for many animals, including *Drosophila* (Helfrich-Förster et al., 2001) and humans (Czeisler et al., 1999). Many humans have increased their exposure to artificial light starting with the invention of electric light, now accelerated by increasing expose to screened devices, thus leading to misalignment of circadian clocks to the external environment (Stothard et al., 2017). In adolescents, this weekly weekday-to-weekend light shifts in sleep-wake cycles are chronic, and prevalent, with light shifts lasting from 3 hours or more (Crowley et al., 2007).

Light shift conditions that closely approximate WLS lead to persistent desynchrony in most of the adult *Drosophila* circadian neural network measured at the single-cell resolution for over a simulated week *ex vivo* strongly coincides with the disruption of circadian regulated behavioral outputs *in vivo*. Adult flies exposed to WLS exhibit transient defects in memory, learning, and sleep stability between 5-6 days of the week. This suggests that for weekly repeated WLS, functional consequences downstream to circadian desynchrony are present during most days of the week. The detriments of clock disruption through light shifts may underlie more severe complications due to cumulative weekly repetitions that may span throughout an individual’s life. Based on the many molecular and circuit-circuit organizational similarities between *Drosophila* and mammals, the circadian neural network responses we measure to weekend light shift conditions may be generalizable to humans and other animals.

## Author contributions

TCH, CN, LR, and PJS designed the research, CN, LR, PH, JDE, TCV, THN, and PJS performed research, CN, LR, TLL, PJS, PH, and NP analyzed data, and TCH, CN and LR wrote the paper.

## Dedication

We dedicate this paper to Michael Buchin who designed the highly sensitive camera at Stanford Photonics that made this work possible. Mr. Buchin passed away on December 19, 2014.

## Acknowledgements

This work was funded by NIH R35 GM127102. We thank Dr. Ralf Stanewsky for the XLG-Per-Luc transgenic fly lines used in the bioluminescence and behavioral experiments. We thank Dr. Lisa Baik for help with designing and editing figures, and manuscript input, Parrish J. Powell and the members of Holmes lab for general laboratory maintenance. We thank Dr. William Joiner (University of California, San Diego) with assistance in sleep analysis, Camden Jansen (Mortazavi Lab) and Lili Li (Wunderlich Lab) for MATLAB assistance.

## Data Availability

All data, MATLAB and R Scripts are available upon request from the corresponding author.

## Extended Data Figure Legends

**Figure S1.**
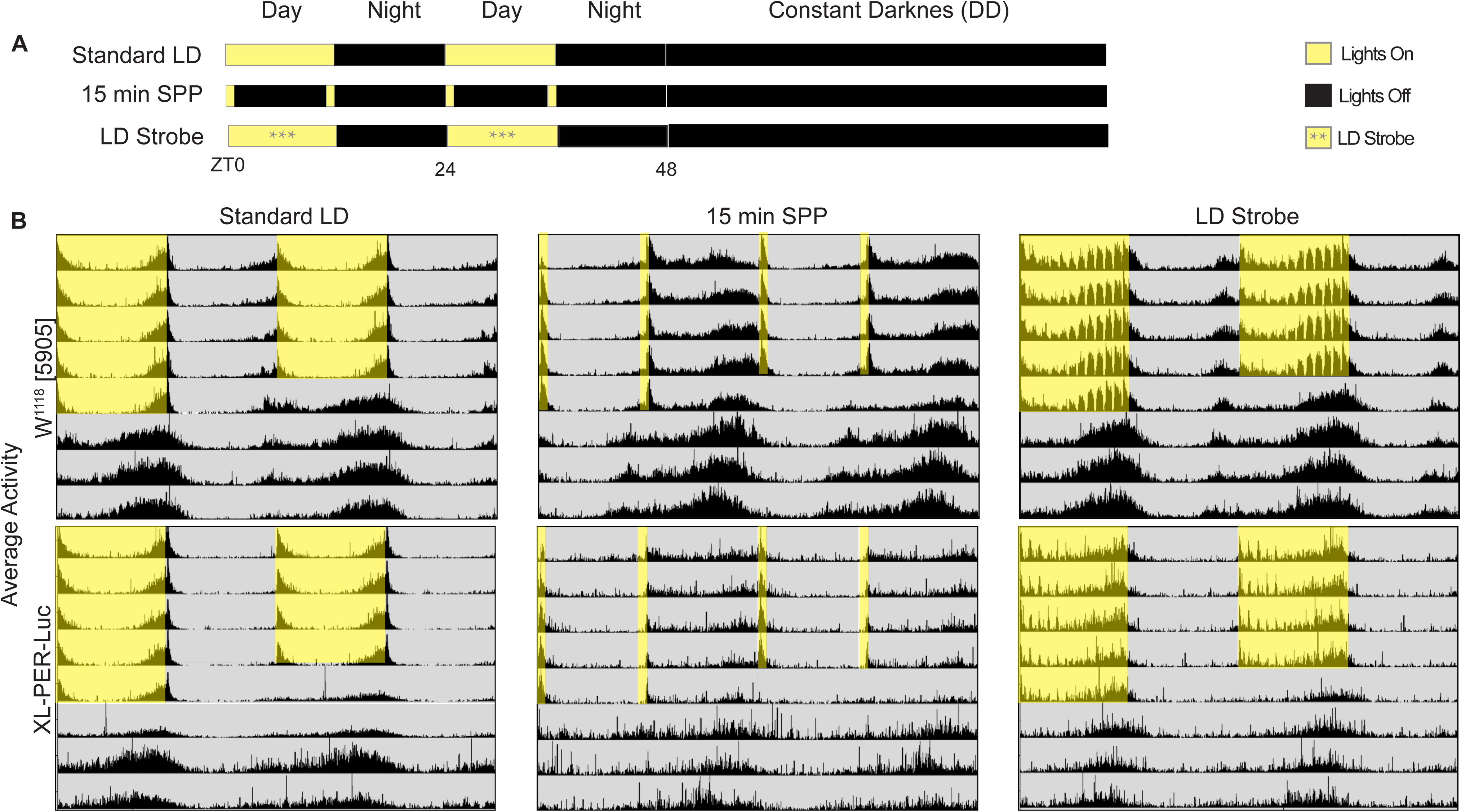
Day-night entrainment of locomotor activity by LD strobe and skeleton photoperiod light protocols. Averaged locomotor activities of adult *Drosophila* for 5 days of entrainment by various light protocols followed by 3 days of constant darkness (DD). All flies were first entrained to ≥3 days of 12:12 hour LD entrainment followed by exposure to LD Strobe (A, top row), Standard 12L12D (A, middle row), or skeleton photoperiod (A, bottom row). Yellow shading indicates intervals of light exposure whereas gray shading indicates intervals of darkness. XLG-Luc flies (B, top row) and wild-type W1118[5905] (B, bottom row) were exposed to 5 days of Standard LD entrainment followed by constant darkness. XLG-Luc flies (C, top row) and wild-type W1118[5905] (C, bottom row) were exposed to 5 days of 15-min skeleton photoperiod followed by constant darkness. XLG-Luc flies (D, top row) and wild-type W1118[5905] (D, bottom row) flies were exposed to 5 days of LD Strobe entrainment followed by constant darkness.

**Figure S2.**
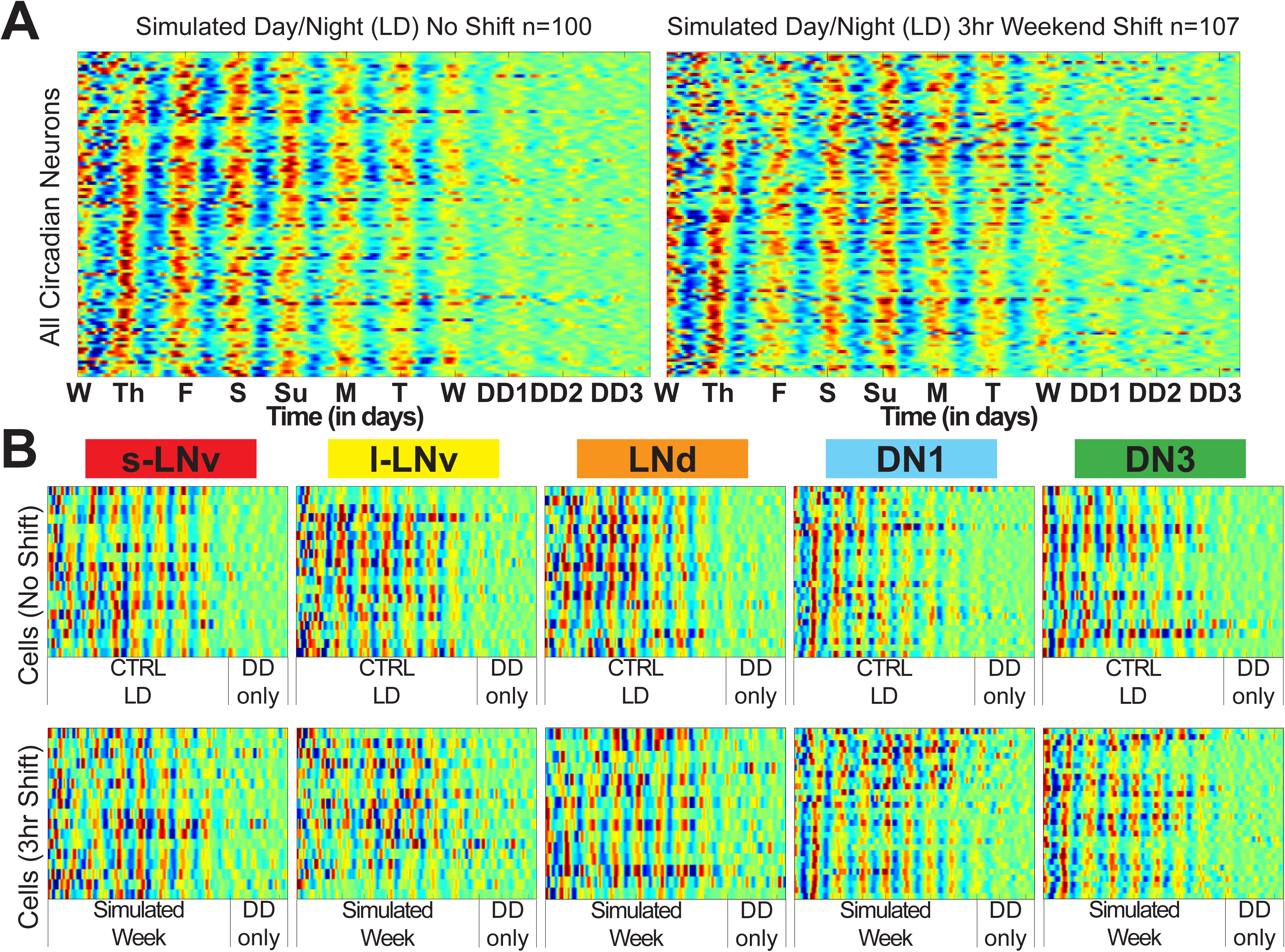
Weekend light shifts lead to circadian neuron desynchrony after one simulated week. Custom MATLAB scripts were employed to create raster plots comparing control LD and WLS conditions. Bioluminescence comparisons between canonical circadian neuron subgroups that undergo control LD (top panels) and WLS conditions (lower panels). Brains are exposed to a total of 8 days in either control LD or WLS conditions, followed by 3 days in complete darkness. Color coding reflects the amplitude in bioluminescence (red=high amplitude, blue=low amplitude).

**Figure S3.**
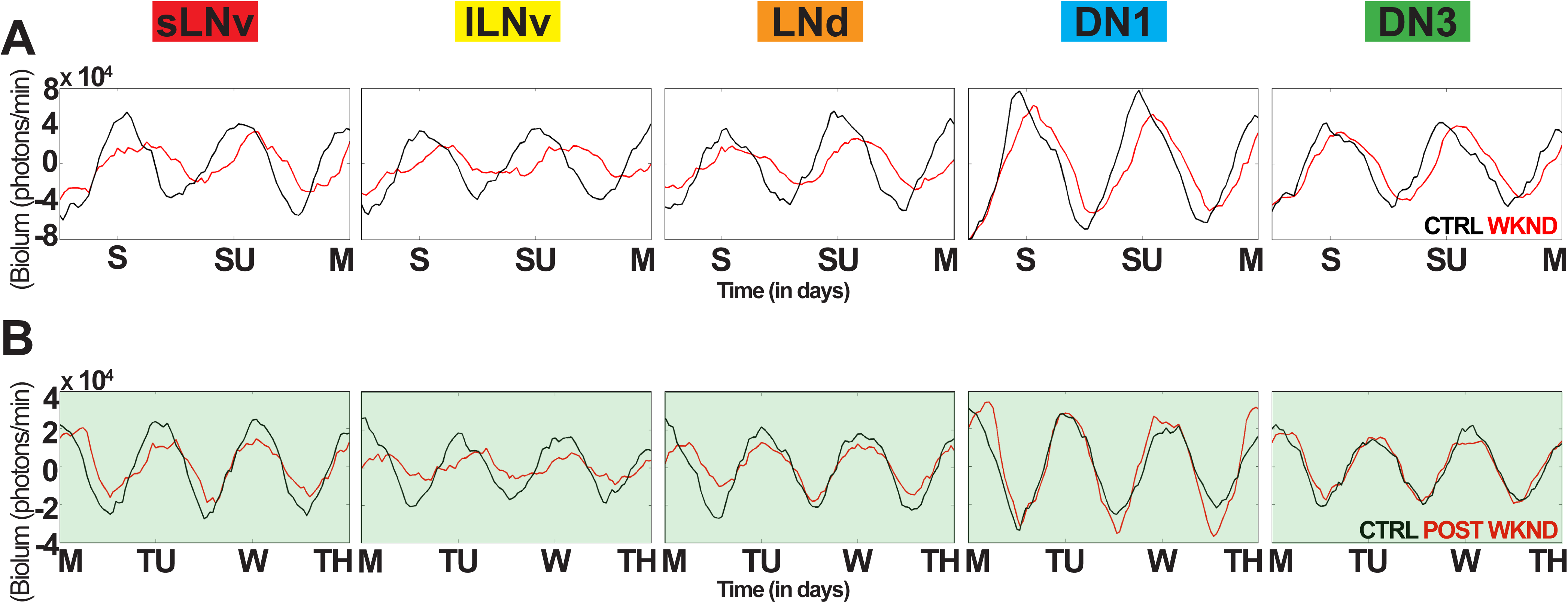
Details of averaged bioluminescent traces during and after 3hr weekend light shifts. (A) Averaged bioluminescence traces for each circadian subgroup comparing the “simulated weekends” in control LD (black trace) with WLS conditions (red trace). (B) Averaged bioluminescence traces for each circadian subgroup comparing the “post-WLS” weekdays in control LD (black trace) and WLS conditions (red trace). Traces for both control LD and WLS conditions were generated using custom MATLAB scripts. See Methods and Materials for more details.

**Figure S4.**
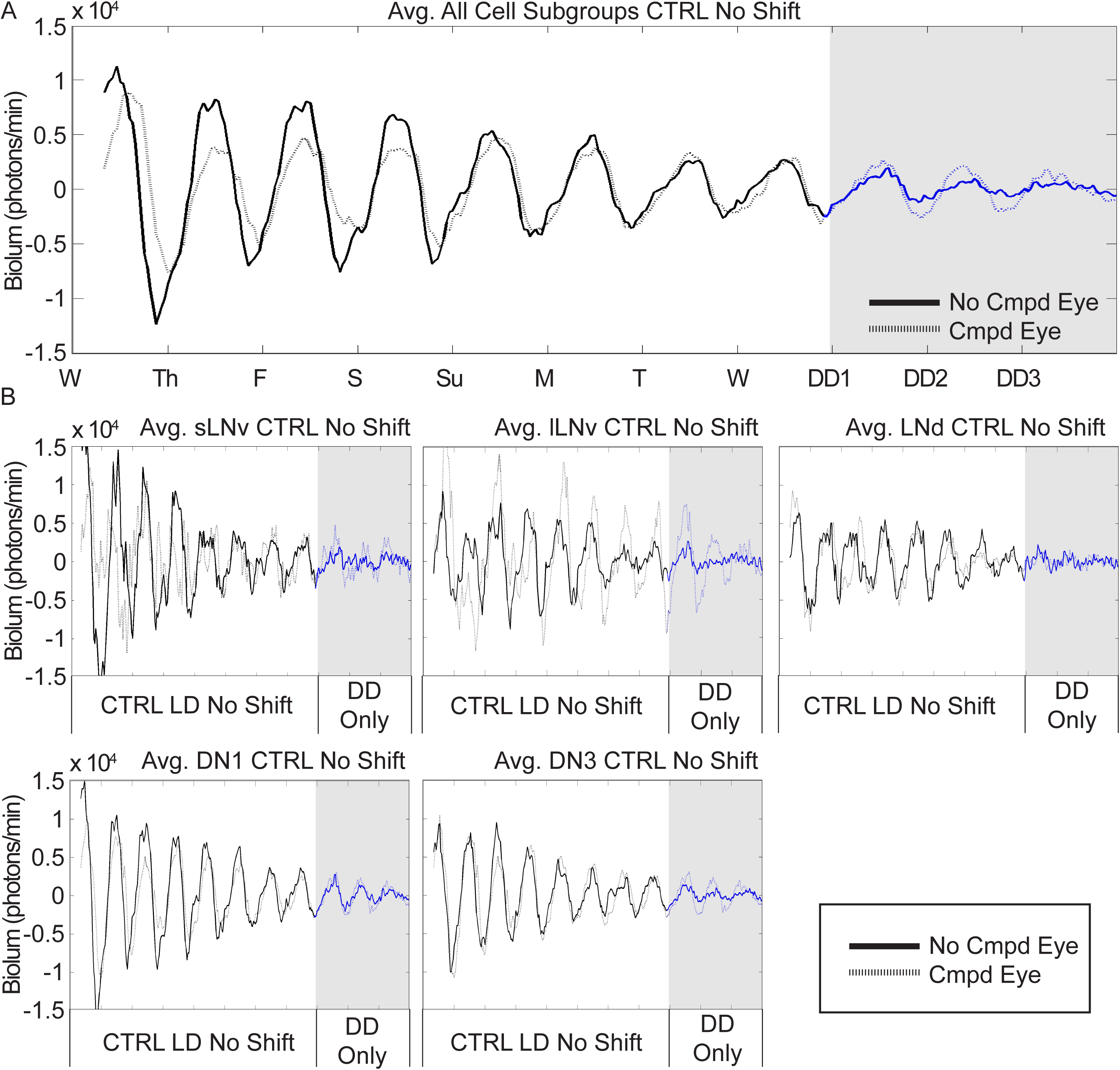
Circadian neuron subgroups are light entrained with or without compound eyes. Averaged 11-day bioluminescence recordings of cultured *Drosophila* brains with compound eyes attached (n=3 brains, dotted plots) or completely removed (n=3 brains, solid plots) reported by PER. (A) Control LD conditions simulate standard 12L12D entrainment spanning one week without phase shifts for cultured brains with compound eyes (sLNv n=6, lLNv n=5, LNd n=7, DN1 n=30, DN3 n=22) and brains with compound eyes removed (sLNv n=7, lLNv n=7, LNd n=7, DN1 n=29, DN3 n=13) followed by DD (gray shade). (B) Averaged bioluminescence traces for individual neuronal subgroups comparing neurons in brains with or without compound eyes.

**Figure S5.**
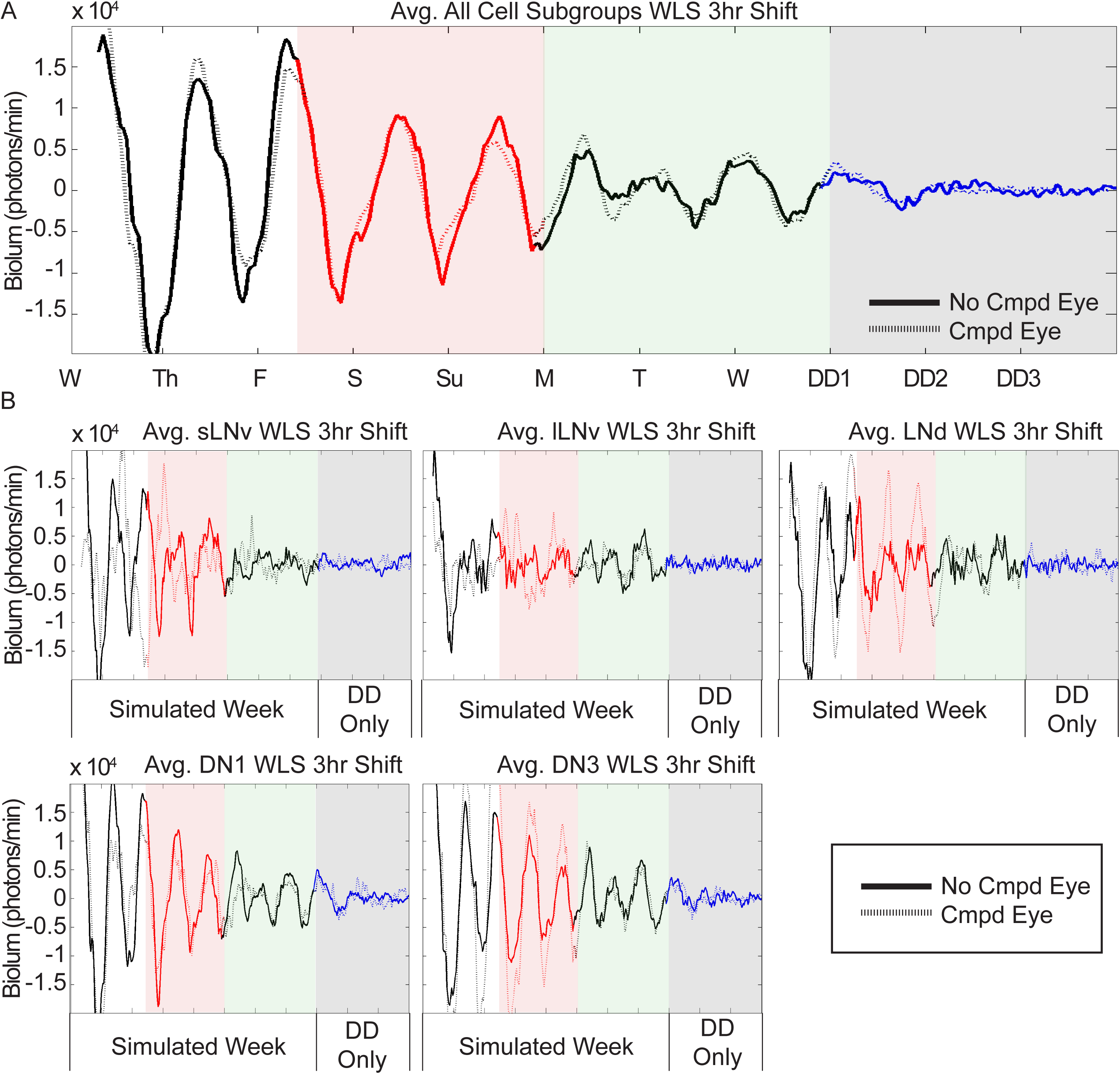
Circadian neuron subgroups respond to phase advances and delays with or without compound eyes. Averaged 11-day bioluminescence recordings of cultured *Drosophila* brains with compound eyes attached (n=3 brains, dotted plots) or completely removed (n=3 brains, solid plots) reported by PER. (A) WLS conditions for each circadian neuron subgroup, each line represents an individual cell in pre-WLS (black trace), WLS (red trace), post-WLS (green shade), then three days of DD (blue trace, gray shade) for brains with compound eyes (sLNv n=5, lLNv n=4, LNd n=3, DN1 n=21, DN3 n=13) and brains with compound eyes removed (sLNv n=4, lLNv n=5, LNd n=7, DN1 n=9, DN3 n=17) followed by DD (gray shade). (B) Averaged bioluminescence traces for individual neuronal subgroups comparing neurons in brains with or without compound eyes under a WLS schedule.

**Table S1.**
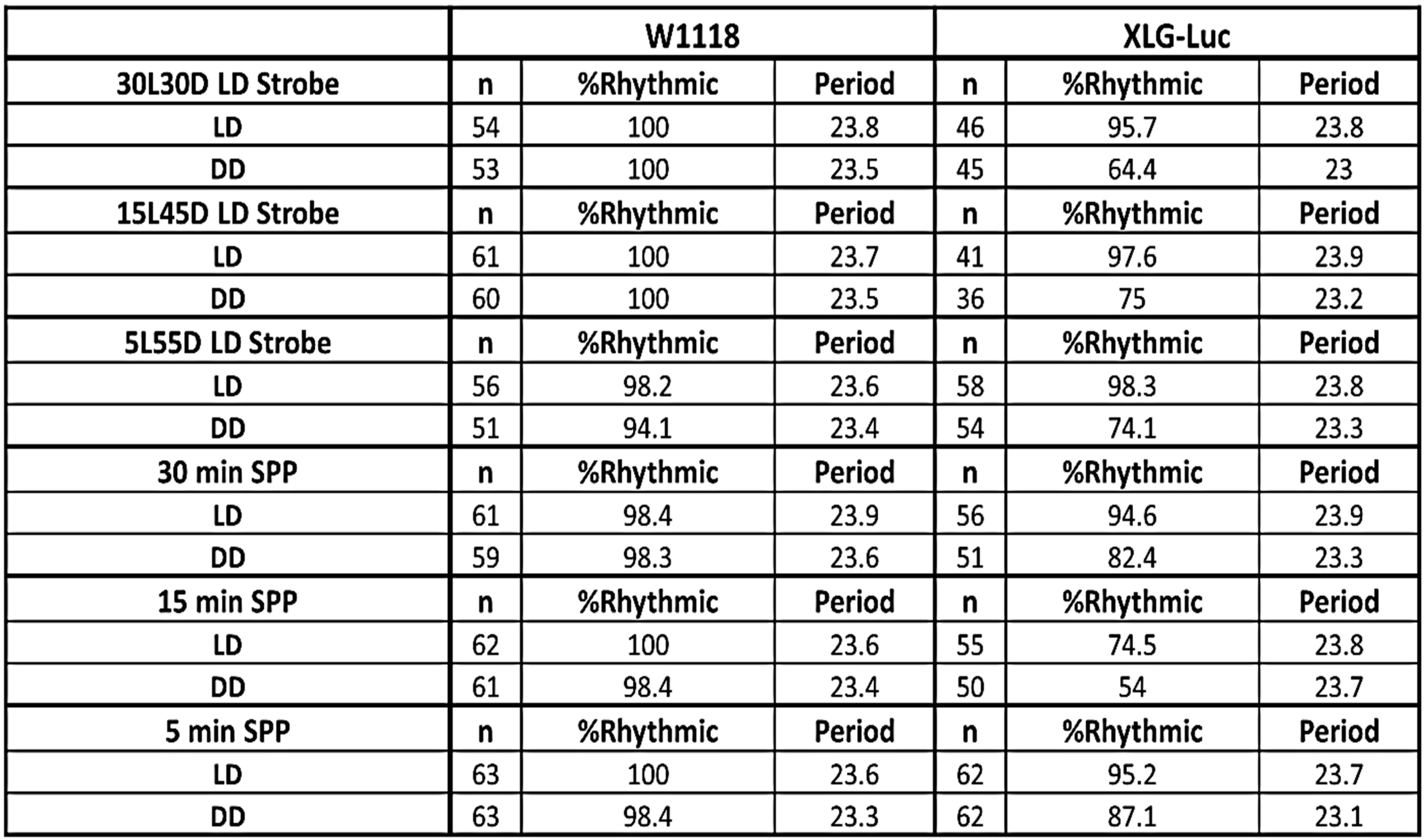
Quantification of behavioral entrainment by LD strobe and skeleton photoperiod. FaasX was used for analysis of behavioral experiments. Wild-type (W1118[5905]) and XLG-luc flies were exposed to 5 days of entrainment (LD) by either LD strobe or skeleton photoperiod protocols with 30, 15 or 5-minute pulses. Following entrainment, flies were maintained in constant darkness (DD) for 5 days. Cycle-P was used to quantify measures of period length and the percentage of rhythmic flies using 15-minute bins of individual fly locomotor activity. Individual flies were considered rhythmic by chi-square periodogram analysis if they met the following criteria: power ≥ 40, width ≥ 4 hours and period length of 24 ± 8 hours.

**Movie S1:**
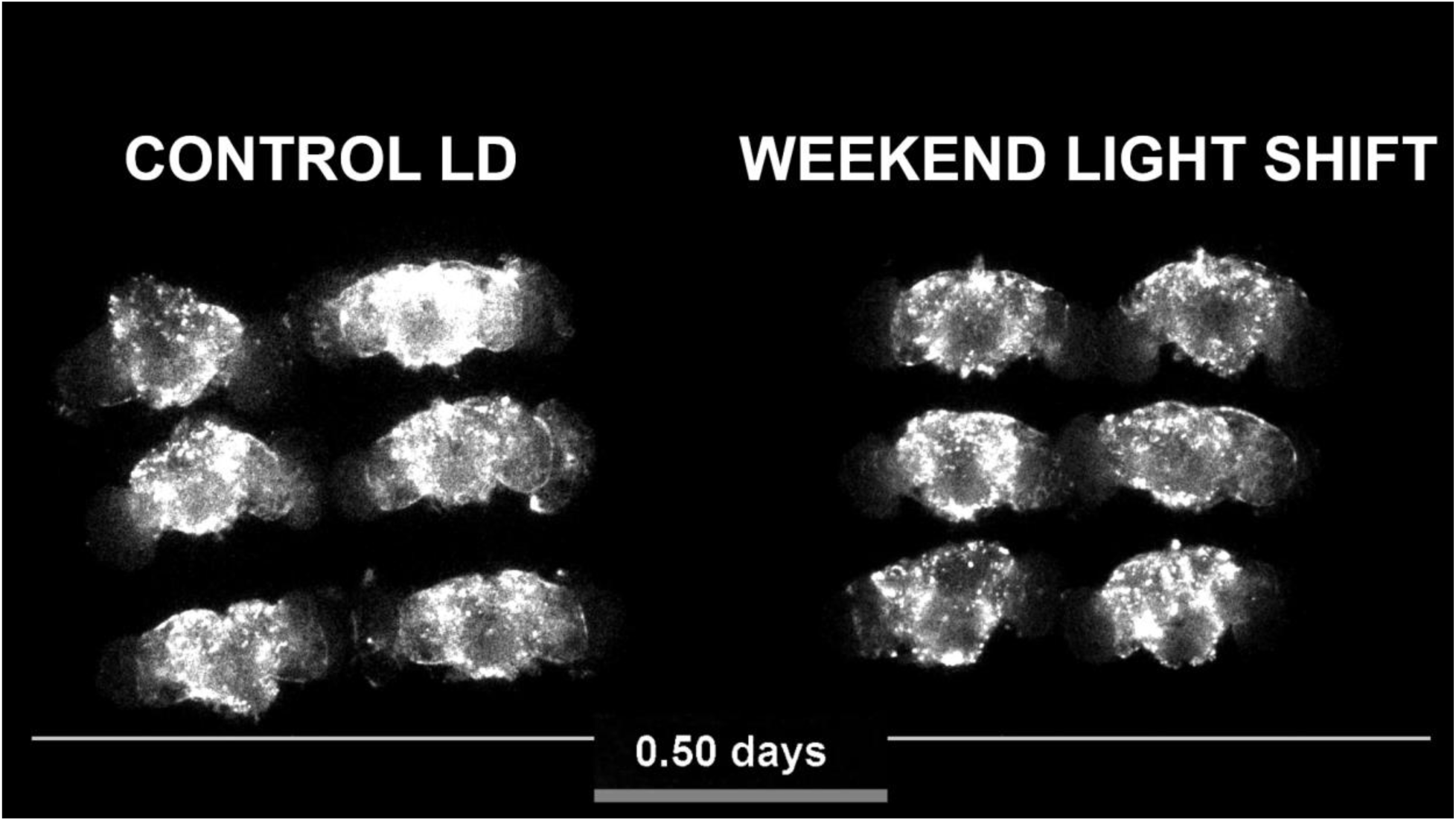
Raw time-lapse bioluminescence recordings of adult XLG-Per-Luc *Drosophila* whole-brain explants cultured for 11 days. Left: Six whole brain culture explants maintained in control conditions (LD strobe with no phase shift) for 9 days followed by 2 days of constant darkness (DD). B: Six whole brain culture explants exposed to weekend light shifts for 9 days followed by transfer to DD for the final 2 days. See Materials and Methods for more details.

**Movie S2:**
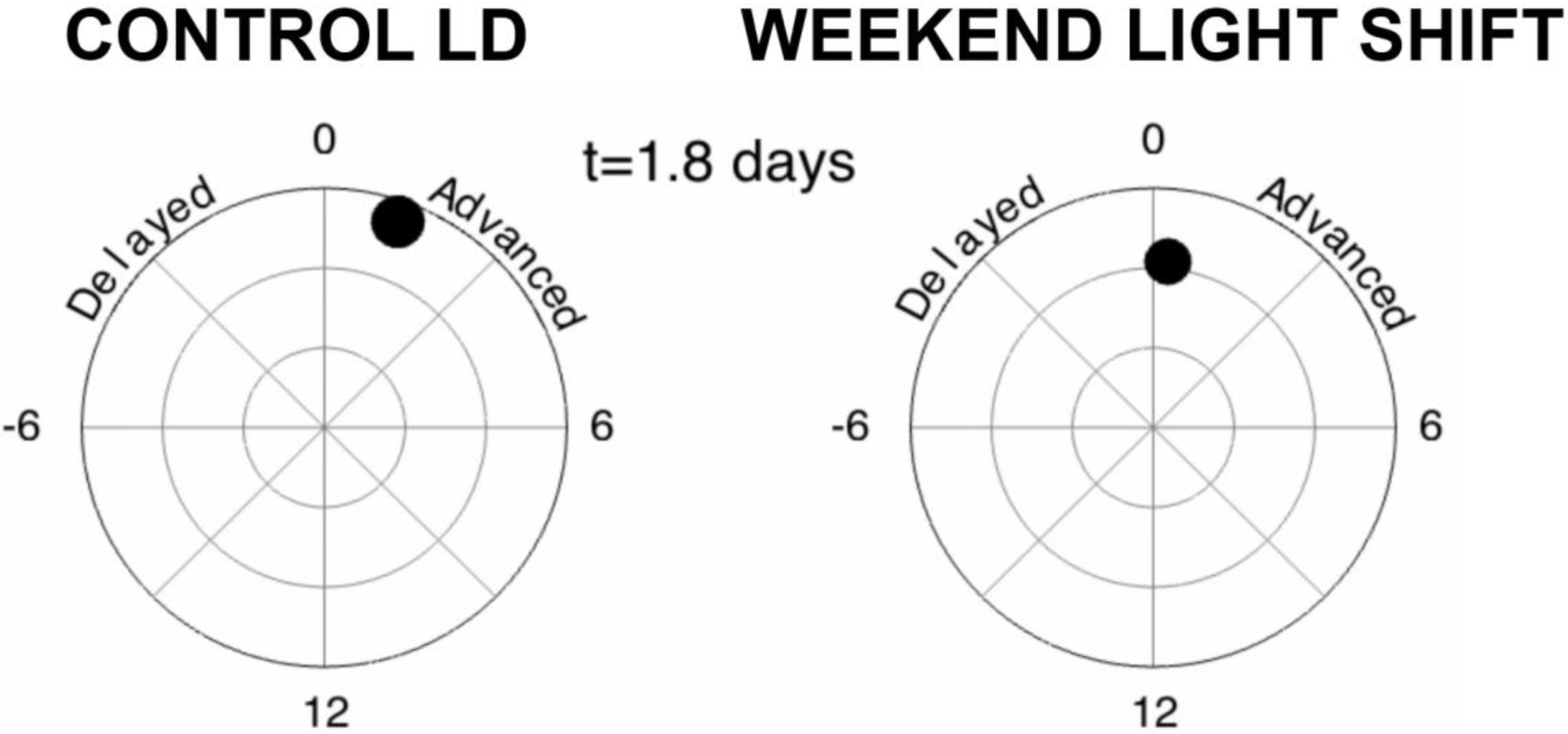
The animations show changes in the relative phase and amplitude of XLG-Per-Luc bioluminescence activity for “all cells” (averaged from all neuronal subgroups) in either control (left) or WLS (right) conditions. Mean network phase is standardized so that the mean network phase is set to ZT 0 on Day 3 when entrainment is most stable. The angle of the disks represents the relative phase shift over time such counterclockwise movement indicates a phase delay whereas clockwise movement indicates a phase advance. The drift of the disks towards the center of the circle and the size of the disks indicates reduction in amplitude. See Materials and Methods for more details.

**Movie S3:**
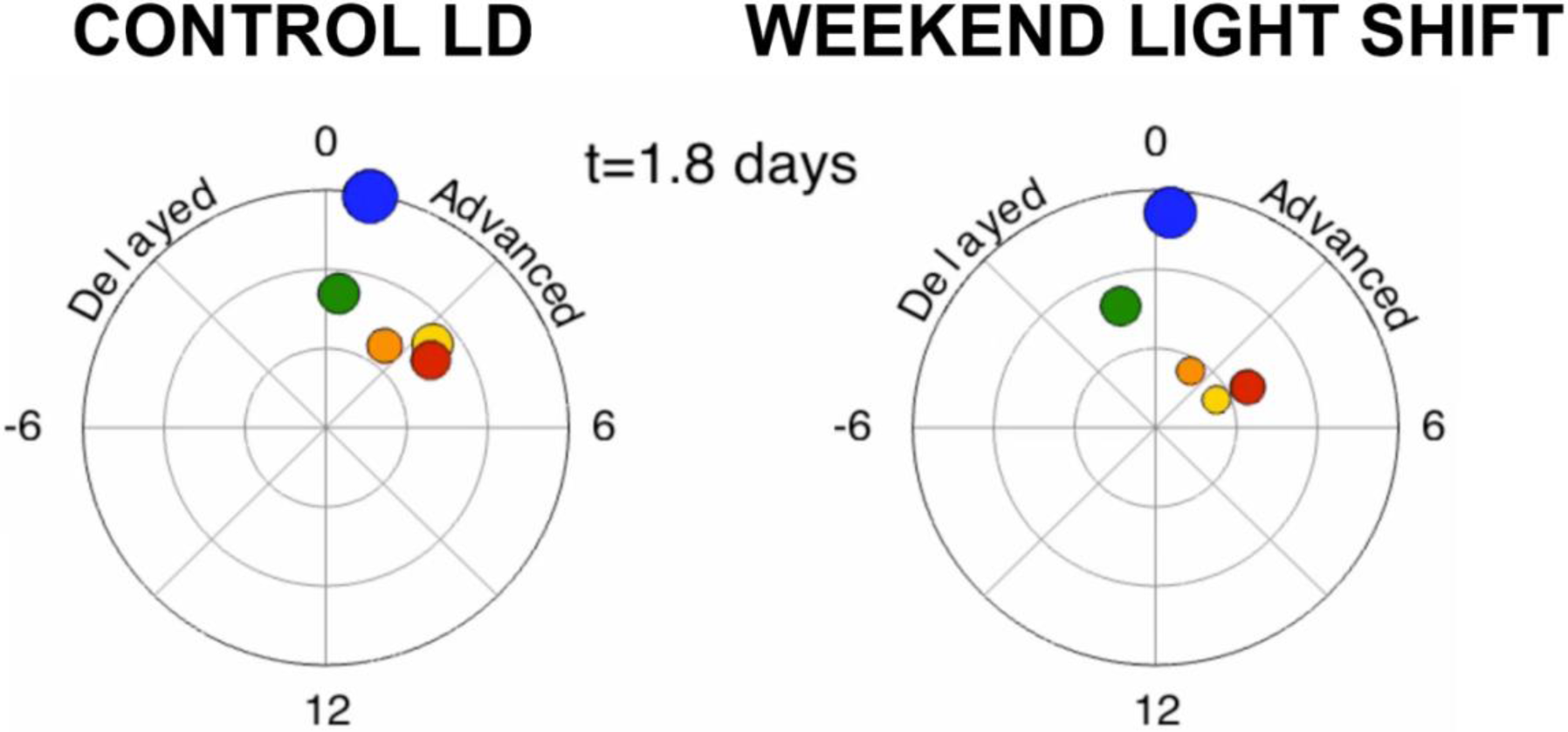
The animations show dynamic changes in relative phase shifts and amplitude for each neuronal subgroup in either control conditions (left) or in response to WLS (right). The mean phase shift for each neuron subgroup is represented by polar angle of the disks whereas amplitude is represented by the size and proximal distance of the disks from the center of the circles. The disks are colored according to neuronal subgroup for the s-LNvs (red), l-LNvs (yellow), LNds (orange), DN1s (blue) and DN3s (green).

**Movie S4:**
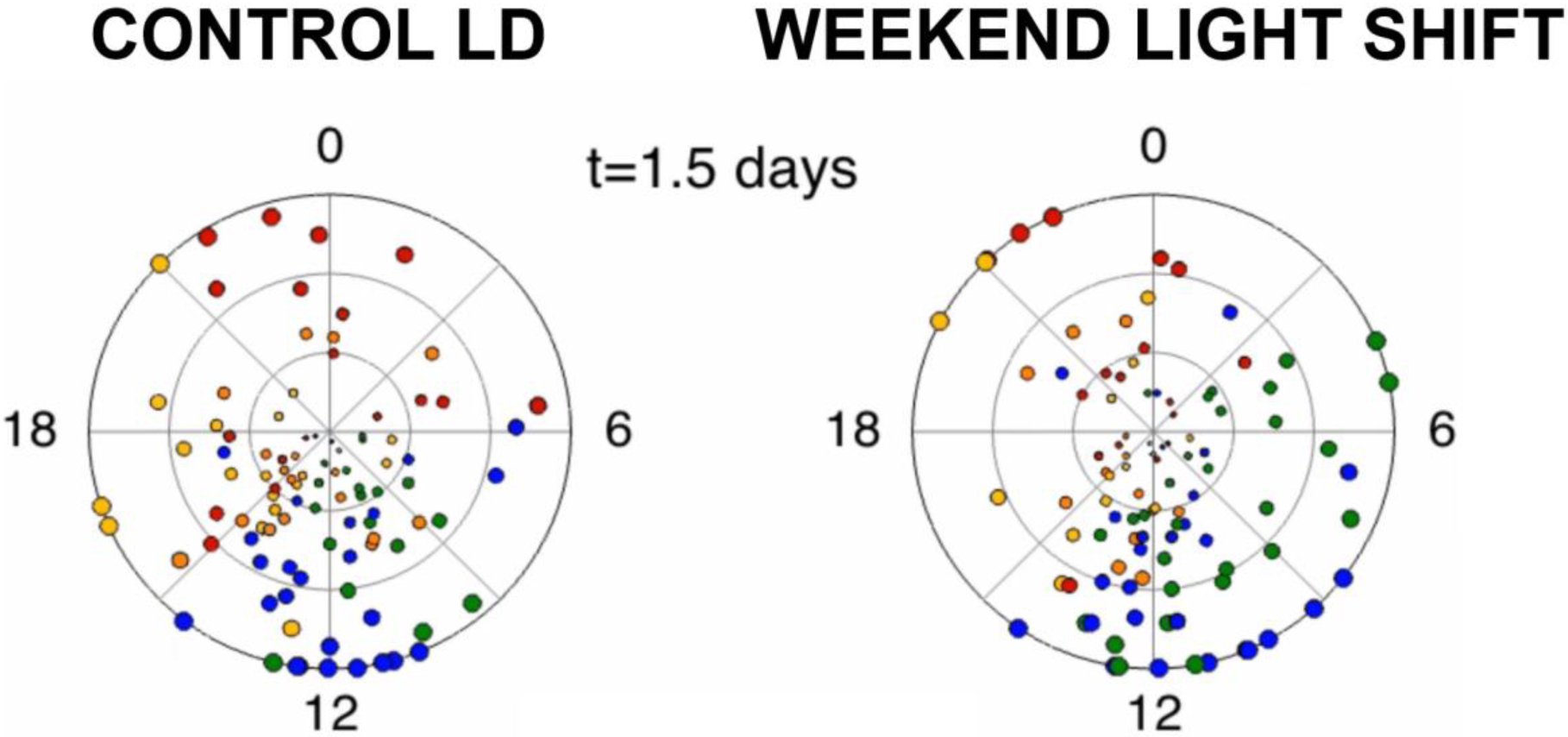
The animations show changes in the phase and amplitude of XLG-Per-Luc bioluminescence activity for individual neuron oscillators from all neuronal subgroups in either control conditions (Left) or in response to WLS (Right). The angle of the disks represents oscillator phase and drift of the disks towards the center of the circle and the size of the disks indicates reduction in amplitude. The disks are colored according to neuronal subgroup for the s-LNvs (red), l-LNvs (yellow), LNds (orange), DN1s (blue) and DN3s (green).

**Movie S5:**
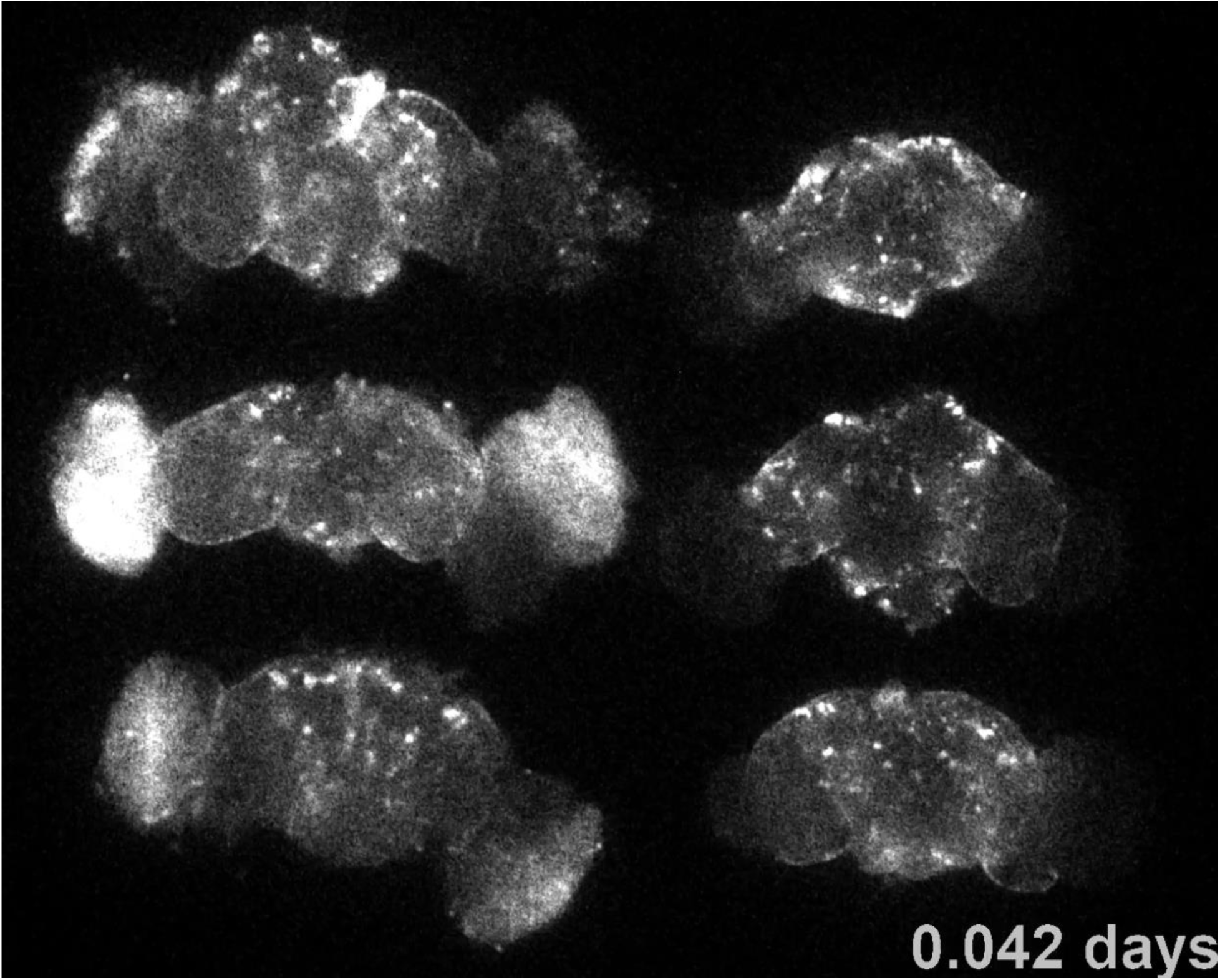
Raw time-lapse recordings of adult XLG-Per-Luc *Drosophila* whole-brain explants comparing bioluminescence of brains with and without compound eyes in CTRL LD. Left: three whole brain culture explants with compound eyes attached maintained in control conditions (LD strobe with no phase shift) for 9 days followed by 2 days of constant darkness (DD). Right: three whole brain culture explants with compound eyes removed maintained in control conditions (LD strobe with no phase shift) for 9 days followed by 2 days of constant darkness (DD). See Materials and Methods for more details.

**Movie S6:**
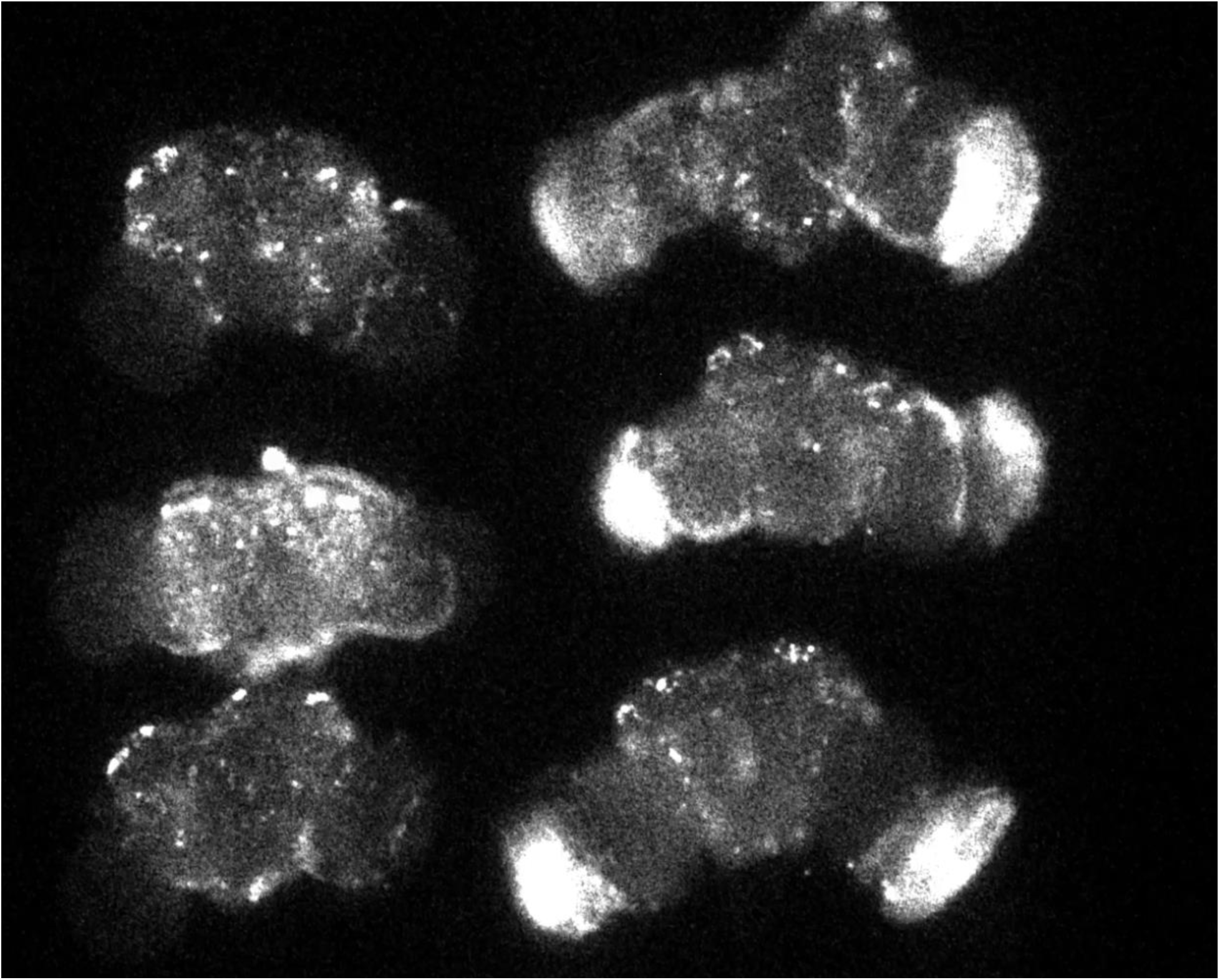
Raw time-lapse recordings of adult XLG-Per-Luc *Drosophila* whole brain explants comparing bioluminescence of brains with and without compound eyes in WLS schedule. Left: three whole brain culture explants with compound eyes removed maintained in WLS conditions (LD strobe with 3hr phase shifts simulating weekends) for 9 days followed by 2 days of constant darkness (DD). Right: three whole brain culture explants with compound eyes attached maintained in WLS conditions (LD strobe with 3hr phase shifts simulating weekends) for 9 days followed by 2 days of constant darkness (DD). See Materials and Methods for more details.

