## Supplementary figures and images for "Weekend light shifts evoke persistent *Drosophila* circadian neural network desynchrony"

### Figure S1

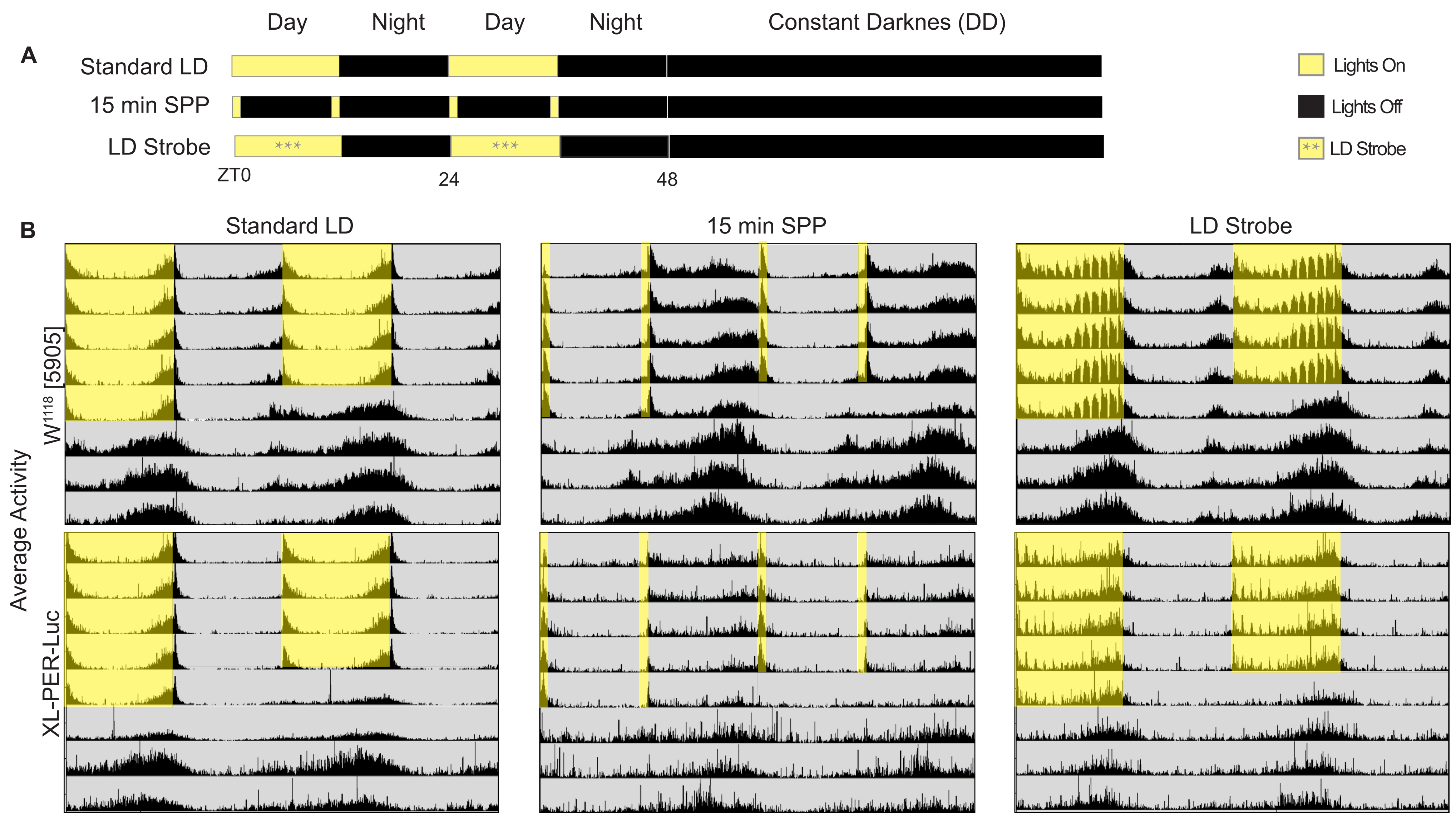

### Figure S2

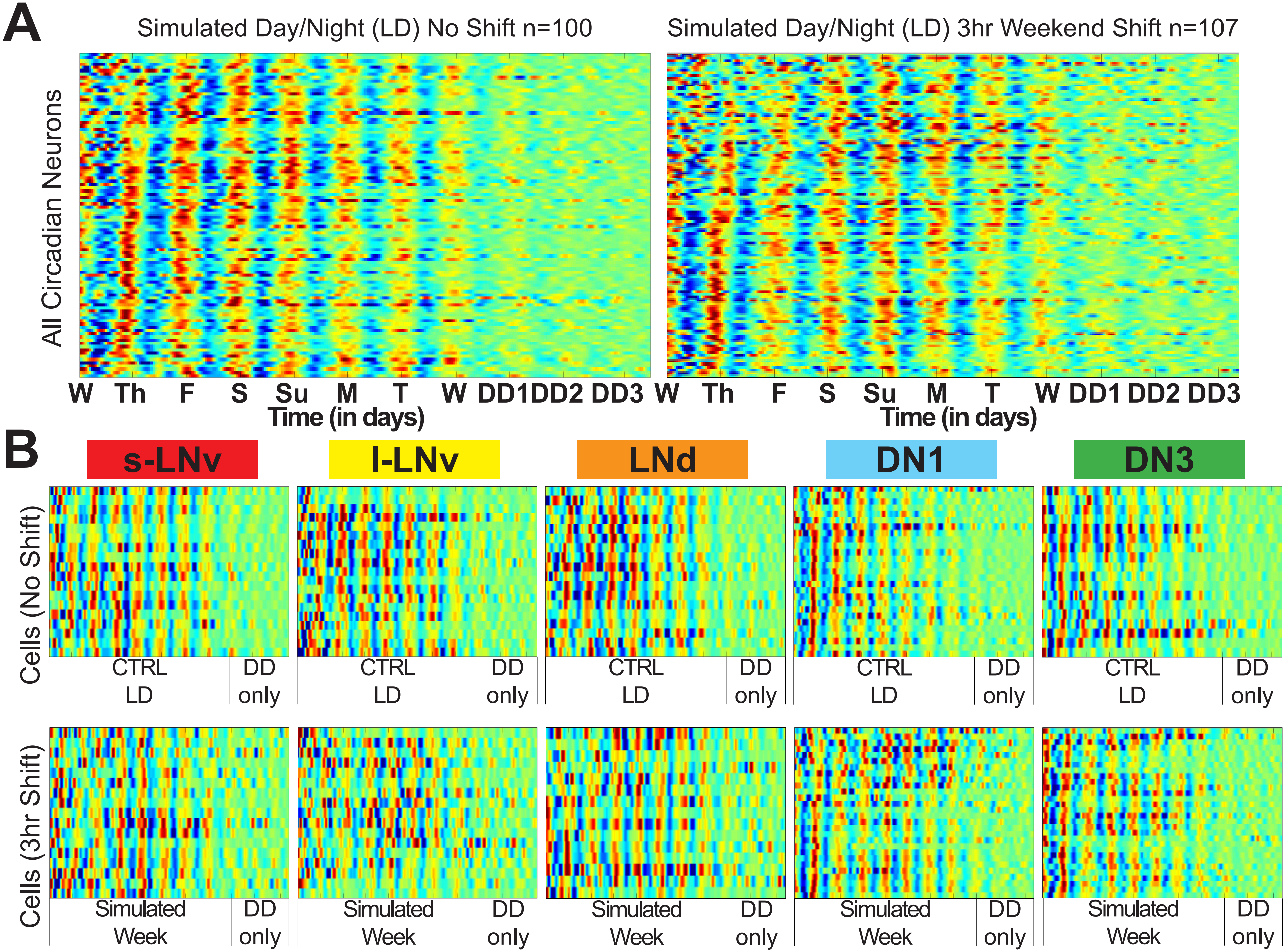

### Figure S3

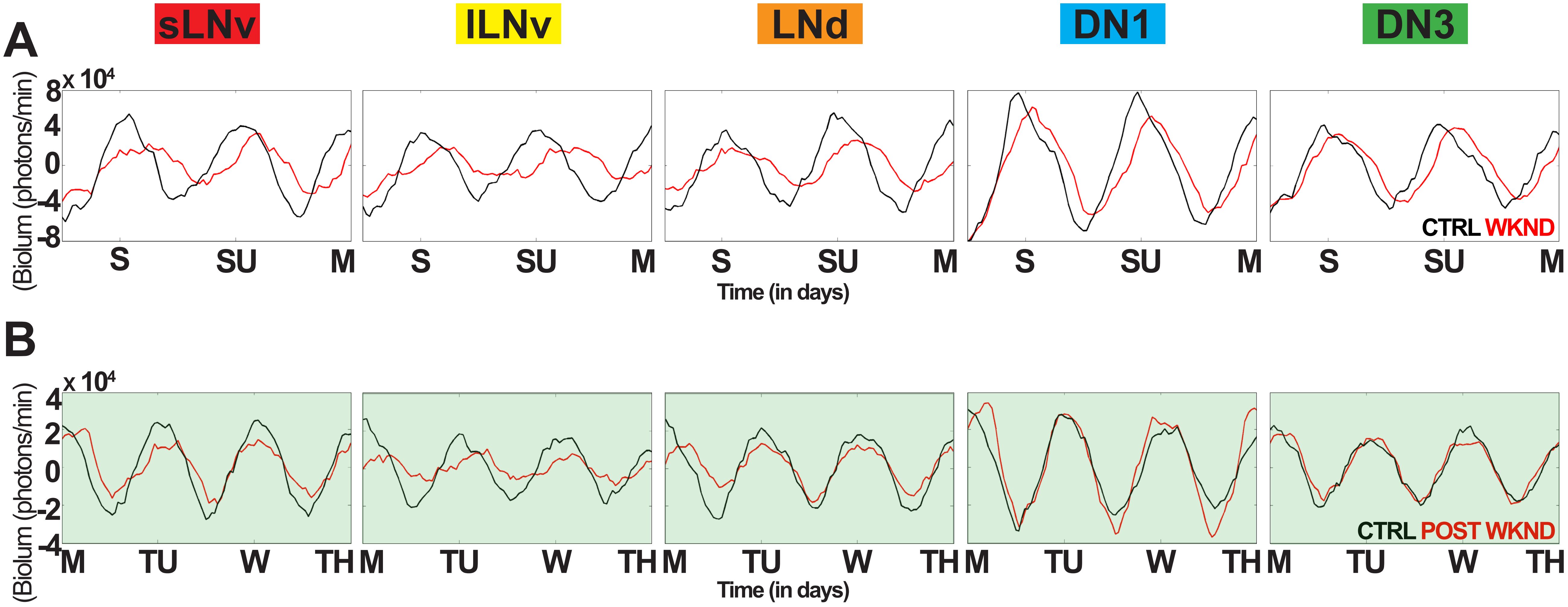

### Figure S4

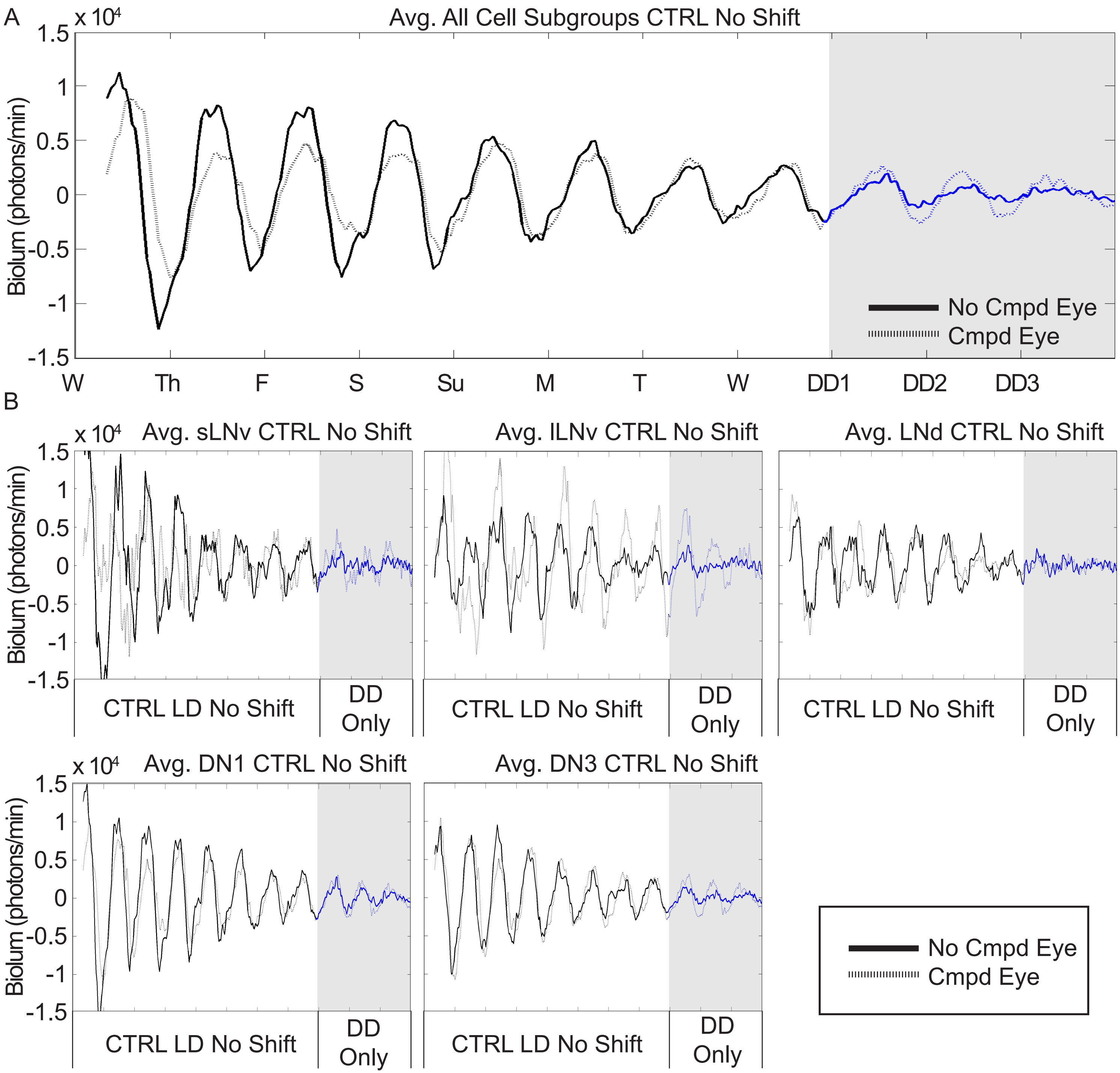

### Figure S5

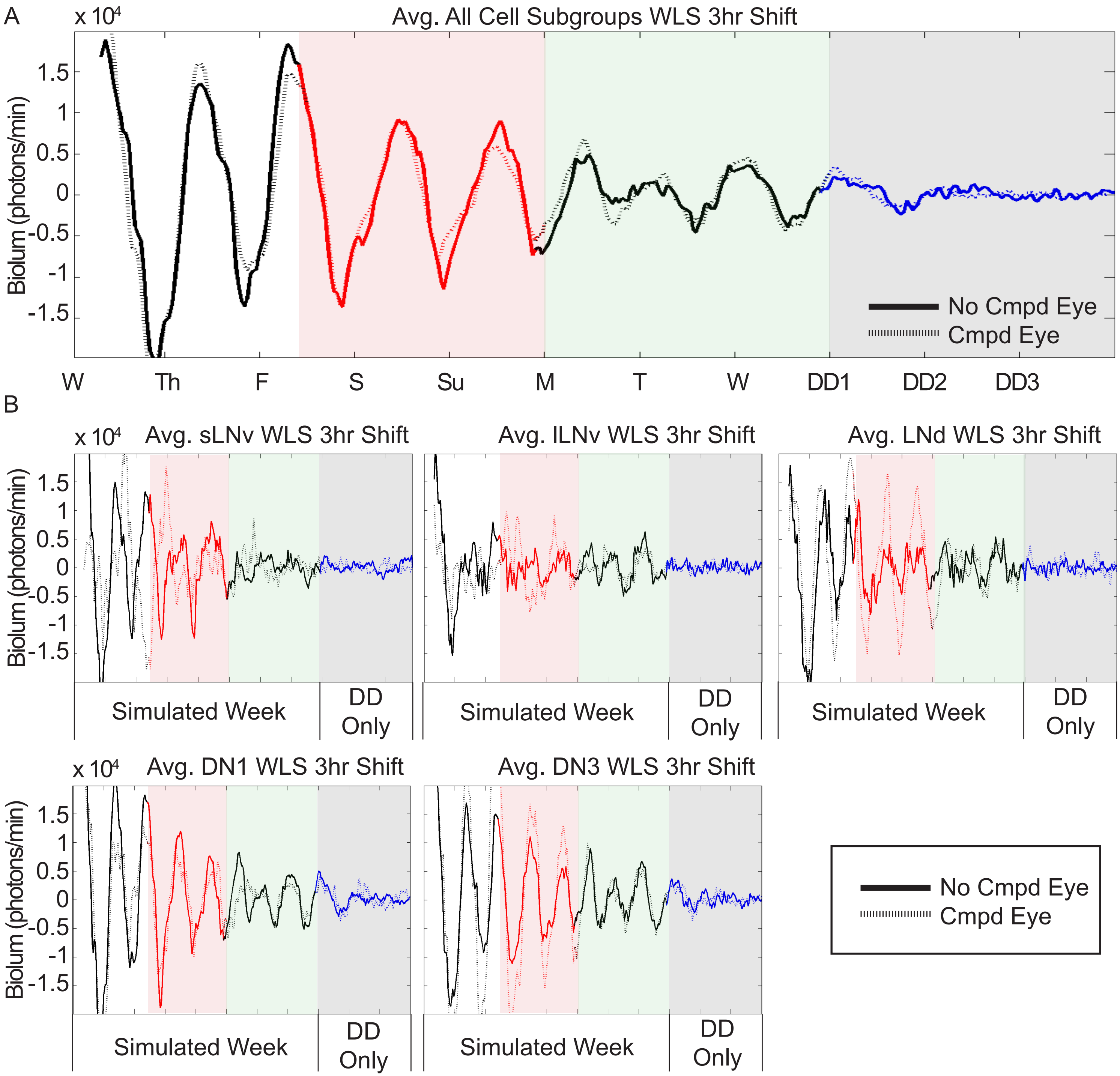
