## Supplementary Table 1 for "Weekend light shifts evoke persistent *Drosophila* circadian neural network desynchrony"

| | W1118 | | | XLG-Luc | | |
|---|---|---|---|---|---|---|
| 30L30D LD Strobe | n | %Rhythmic | Period | n | %Rhythmic | Period |
| LD | 54 | 100 | 23.8 | 46 | 95.7 | 23.8 |
| DD | 53 | 100 | 23.5 | 45 | 64.4 | 23 |
| 15L45D LD Strobe | n | %Rhythmic | Period | n | %Rhythmic | Period |
| LD | 61 | 100 | 23.7 | 41 | 97.6 | 23.9 |
| DD | 60 | 100 | 23.5 | 36 | 75 | 23.2 |
| 5L55D LD Strobe | n | %Rhythmic | Period | n | %Rhythmic | Period |
| LD | 56 | 98.2 | 23.6 | 58 | 98.3 | 23.8 |
| DD | 51 | 94.1 | 23.4 | 54 | 74.1 | 23.3 |
| 30 min SPP | n | %Rhythmic | Period | n | %Rhythmic | Period |
| LD | 61 | 98.4 | 23.9 | 56 | 94.6 | 23.9 |
| DD | 59 | 98.3 | 23.6 | 51 | 82.4 | 23.3 |
| 15 min SPP | n | %Rhythmic | Period | n | %Rhythmic | Period |
| LD | 62 | 100 | 23.6 | 55 | 74.5 | 23.8 |
| DD | 61 | 98.4 | 23.4 | 50 | 54 | 23.7 |
| 5 min SPP | n | %Rhythmic | Period | n | %Rhythmic | Period |
| LD | 63 | 100 | 23.6 | 62 | 95.2 | 23.7 |
| DD | 63 | 98.4 | 23.3 | 62 | 87.1 | 23.1 |
